# Comprehensive engineering of the CasX-Editor to generate a therapeutically optimized genome editor

**DOI:** 10.64898/2026.11.10.765514

**Authors:** Addison V Wright, Sean A Higgins, Elizabeth J Bailey, Ross White, Ashley E Graham, Isabel Colin, Matthew R Gardner, Kristin Wucherer, Cécile Fortuny, Jennifer Saraya, Shyam Sundhar Bale, Santhosh Karanth, TC Steven Keller, Michelle Eggers, Farah H Bardai, Kristina Bender, Myles Hochman, Benjamin Demaree, Shiyi Wang, Kirsten A Reimer, Keith Szulwach, Christie C Sze, Allison Didychuk, Sarah K Denny, Benjamin L Oakes

## Abstract

CRISPR-based genetic modification is enabling curative, one-time treatments, but extending its use to more diseases will require increasing potency at low doses while avoiding unintended genomic modification. CRISPR technologies derive from microbial nucleases, such as Cas9, whose properties evolved for antiviral defense rather than precision editing in human cells. Although these systems can be engineered for greater activity or specificity, improving both properties to meet the challenges of *in vivo* delivery in large animals remains challenging. Here we used iterative molecular engineering of the Cas12e protein and its guide RNA to produce the X-Editor (XE), a compact nuclease platform that combines robust *in vivo* activity with exceptional genomic specificity. Screening tens of thousands of protein variants and guide designs and iteratively testing and combining beneficial mutations yielded an enzyme with more than 100-fold higher editing activity in human cells. Cryo-electron microscopy revealed novel inter- and intramolecular interactions driving enhanced function. XE mRNA was delivered to nonhuman primate liver via lipid nanoparticle with guides for two distinct targets. At both targets, XE demonstrated greater editing at lower doses than comparable preclinical data from Cas9-based therapies that are now in clinical testing. No off-target editing was detected for either guide in full specificity assessments performed in primary human hepatocytes at doses tenfold above the on-target effective concentration (EC90). These findings establish that more potent *in vivo* editing and stringent genomic specificity can be jointly engineered into a naturally compact CRISPR system for delivery using adeno-associated virus (AAV) or lipid nanoparticle (LNP) delivery strategies. These results provide a new foundation for advanced genetic medicines that demand lower dosing and greatly reduced off-target risk.

## INTRODUCTION

Genetic medicine holds the promise of one-time, curative treatments by installing beneficial edits to our DNA. CRISPR-Cas systems have revolutionized the field by making gene editing tractable and flexible via RNA-programmable DNA targeting. Multiple CRISPR-based therapies have entered the clinic, predominantly targeting severe genetic disorders and culminating in the first approved CRISPR-based therapy for sickle cell disease^1–6^. This rapid clinical translation has been driven largely by the remarkable activity of *Streptococcus pyogenes* Cas9 (SpCas9) nuclease, which remains the foundation of most therapeutic genome editing technologies, including base editing and prime editing^7^.

However, extending genome editing beyond severe genetic disorders to common diseases affecting millions of patients demands a substantially wider therapeutic window, in which potent editing can be achieved at low, clinically practical doses while minimizing dose-dependent toxicity and unintended genomic modification. Overcoming this challenge requires more than incremental, isolated improvements in nuclease activity or specificity. Instead, CRISPR systems must be comprehensively engineered to jointly optimize potency and specificity to enable saturating editing at low doses without compromising safety.

Almost all CRISPR-based therapeutic approaches utilize minimally modified nucleases which did not evolve for therapeutic genome editing. As components of prokaryotic immune systems, these enzymes evolved to recognize and destroy invading genetic elements under selective pressures fundamentally different from those governing therapeutic performance in human cells. High specificity is deleterious in anti-phage defense, as phage mutations would readily enable immune evasion. Further, phage and prokaryotic genomes present a fundamentally different substrate than a human genome due to their smaller size and lack of nuclear envelope or chromatinized DNA. Thus, the properties that make a CRISPR nuclease successful in nature are not necessarily those required for a human therapeutic.

Considerable efforts have sought to overcome these limitations through protein and RNA engineering in one-off fashion, introducing improvements in activity, specificity or targeting range to SpCas9 and other Cas effectors^8–16^. However, these properties are rarely improved in tandem, and the efficacy of the resulting molecules is rarely demonstrated under the challenging conditions of *in vivo* delivery in large animal models. More recently, machine learning and generative protein models trained on sequence diversity have expanded the repertoire of CRISPR and related nucleases, producing functional proteins diverged from individual natural effectors^17,18^. While these approaches demonstrate that recombining natural solutions can generate functional sequences, their predictions are limited by sequence landscapes generated by evolutionary pressures not aligned with therapeutic goals. Achieving a fundamentally distinct balance of potency and specificity therefore requires engineering guided by empirical selection for therapeutic performance and systematic improvement over multiple engineering cycles.

We show here that the CasX (Cas12e) ribonucleoprotein (RNP), a compact and structurally distinct CRISPR nuclease with favorable target discrimination but insufficient activity in mammalian cells, can be engineered to enhance activity while maintaining high specificity^19–22^. Coordinated engineering of the CasX protein and its guide RNA, using successive rounds of rational design, high-throughput selection, and empirical characterization, produced the X-Editor (XE). XE has 113 mutations across the protein and guide compared to naturally occurring CasX systems, with over 75-fold enhanced biochemical cleavage activity and more than 100-fold increased editing activity in human cells. Cryo-electron microscopy (Cryo-EM) of the engineered ribonucleoprotein bound to target DNA reveals novel molecular interactions associated with enhanced activity, providing mechanistic insight and a structural foundation for further engineering. Lipid nanoparticle delivery of XE achieved saturating editing at two distinct therapeutic targets in nonhuman primate liver, exceeding 70% editing in bulk liver tissue at therapeutically relevant doses while no off-target editing was detected for either guide in primary human hepatocytes at exposures tenfold above the on-target EC90. Furthermore, XE demonstrates highly potent, delivery-limited genome editing *in vivo* in multiple tissues when delivered via AAV. This work demonstrates how comprehensive molecular engineering can overcome the functional limitations of naturally evolved CRISPR systems to create precise genome editors designed for the demands of *in vivo* human medicine.

## RESULTS

### Rational engineering and directed evolution of the guide RNA scaffold improves activity

Multiple CasX enzymes have been characterized to date, but none have been shown to edit mammalian genomes efficiently^20,22,23^. We selected PlmCasX as the starting enzyme because it previously showed higher mammalian cell editing activity than DpbCasX^20,22^. We initially focused on engineering the CasX guide scaffold to improve editing efficiency.

We first designed rational modifications to the PlmCasX sgRNA. Our analysis of RNAseq data from the initial description of CasX suggested that transcribed guide *in vivo* includes additional nucleotides 5′ of the terminus compared to those used in subsequent structural and functional characterizations^19,20,22^. We tested variants with additional nucleotides from the native locus at the 5′ end and found that a three-nucleotide extension had the best overall activity (Fig. S1). This 5′ end was used in all subsequent guide designs. Several alternate extended stems were tested to replace the cr:tracr duplex, and truncated versions were found to generally improve editing (Fig. S1). Finally, a homopolymeric stretch of U residues early in the scaffold was disrupted by substituting the first U to a C to avoid premature transcription termination (as in Chen et al, 2014)^24^. The first U in the tract was selected for mutagenesis to avoid disrupting the triplex motif. While this substitution reduced activity (Fig. S1), we reasoned that this variant might serve as a better base for additional engineering due to its higher anticipated transcription. We combined the modifications to the 5′ end, the extended stem, and the polyU stretch into a new gRNA scaffold, called Sv1 (Fig. 1A).

**Figure 1.**
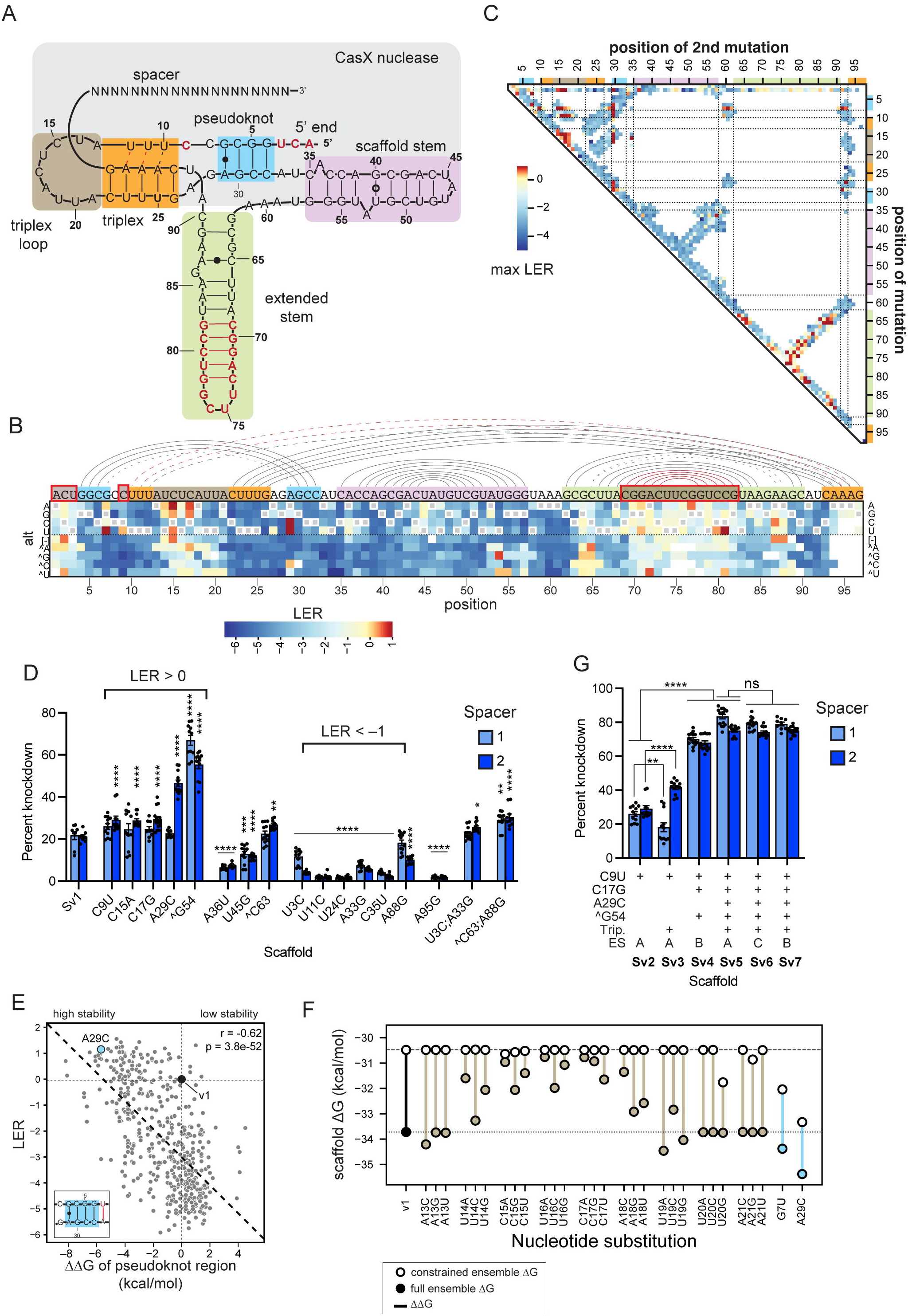
Pooled screening and structure-guided engineering of the CasX sgRNA enable large gains in activity. (A) Structure and domains of the PlmCasX-derived sgRNA Sv1. Bases in red are differences from the PlmCasX version in the 7WB1 cryo-EM structure (Tsuchida et al). (B) Single mutagenesis heatmap shows the Log2 enrichment relative to Sv1 (LER) for all tested single mutations across the Sv1 scaffold. Grey circles indicate the wildtype sequence. Insertions or deletions that are redundant with another pixel within the heatmap have values plotted in both locations. The lines at top indicate basepairing interactions between residues in the 7WB1 structure (or inferred from for mutated sequence - red). Colors indicate the domains as in (A). (C) Double mutagenesis heatmap shows LER for all tested double mutations across the Sv1 scaffold, where the maximum LER across all double mutations that address that pair of positions is shown for each pair of positions. Dotted lines within the double mutagenesis heatmap indicate boundaries of secondary structure regions shown in (A). (D) Comparison of activity of sgRNA variants on HLA knockdown in HEK293T cells with two *B2M*-targeting spacers when delivered via plasmid lipofection with CasX. Mean, standard deviation, and individual points for all replicates meeting inclusion criteria are shown. p-values relative to the corresponding spacer for Sv1 are shown for each bar (one-way ANOVA, Dunnett’s multiple comparison test). * = p<0.05; ** = p<0.01; *** = p<0.001; **** = p<0.0001. (E) Correlation of the pseudoknot stem folding stability and the LER of the sgRNA variant containing that pseudoknot stem sequence. Relative stability (ΔΔG) is obtained by taking the difference between the calculated stability of each variant and that of the Sv1 pseudoknot sequence. The Pearson r value and significance of the linear relationship are indicated on the graph. (F) Quantification of the impact of triplex loop and other mutations on the difference in stability (ΔΔG) between the full predicted folding ensemble and the ensemble expected to be capable of RNP binding. (G) Comparison of activity of additional sgRNA variants on HLA knockdown in HEK293T cells. Mean, standard deviation, and individual points for all replicates meeting inclusion criteria are shown. p-values relative to the corresponding spacer for the indicated comparison are shown (one-way ANOVA, Dunnett’s multiple comparison test). ** = p<0.01; **** = p<0.0001; ns = not significant (p>0.05).

To identify beneficial mutations in an unbiased manner, we designed and screened a library of ∼20,000 scaffold variants comprising all possible single substitutions, insertions, and deletions, as well as double mutations in potentially interacting residues and higher order mutations within defined structural elements. The library was screened in human cells with a spacer targeting a fluorescent reporter to enable enrichment of variants with improved editing (Fig. S2).

Single mutations were assessed for enrichment (Fig. 1B). We found 7.5% of single mutations that had positive Log2 Enrichment relative to Reference (LER). Mutations with positive LER were identified across all domains, with multiple beneficial mutations at many unpaired residues (Fig. 1B). Outside of the extended stem (discussed below), mutations at sites not involved in base-pairing were more than twice as likely to result in positive LER (odds ratio = 2.3). The double mutation data had a similar pattern as the singles (Fig. 1C), though only 3% of double mutations had positive LER. We identified examples of positive epistasis, where double mutations were enriched where no single mutation at either position was a hit, including a base pair generated by our 5′ extension (U3:A33; Fig. S3).

To validate the screen results, we designed variants and tested them with two spacers that target the *B2M* gene, which is essential for HLA presentation, and assessed cells by flow cytometry for HLA knockdown. We selected mutations with positive, slightly negative (>-1), or strongly negative (<-1) LER. We found strong correlation with the screen enrichment data, with all mutations with strongly negative LER having significantly lower activity than the Sv1 scaffold for both spacers, and each of the positive LER variants having improved activity for at least one spacer tested gene (Fig. 1D). To test the prediction of positive epistasis, we tested the combined mutation of U3 and A33 (U3C:A33G) as well as the negatively enriched individual mutations and saw that the double mutant was able to rescue the reduced activity observed with each of the single mutations (Fig. 1D). Overall, the correlation of activity between the screen enrichment and the knockdown measurements supports that the screen provides a quantitative sequence-function map of the gRNA.

We next investigated the relationship between predicted folding stability and activity. Multiple mutations to the noncanonical G7:A29 pair in the pseudoknot stem had positive LER, and A29C, which resolves the pair to a G:C, showed a ∼2-fold improvement in activity for one of the spacers in validation testing (Fig. 1B, 1D). We investigated a set of higher-order mutations included in the library that heavily mutagenized the pseudoknot stem while maintaining predicted structure. LER values were compared to predicted thermodynamic stability (ΔG), and a significant negative correlation was identified (r = –0.62, p <1x10^−50^, Fig. 1E), indicating that more stable pseudoknot sequences support higher guide activity. Notably, Sv1 demonstrated high enrichment relative to pseudoknots with similar stability, which highlights that secondary structure stability alone is not sufficient for optimal activity.

The triplex loop, particularly positions 14-17, showed high tolerance to mutation (Fig. 1B). Secondary structure predictions revealed a potential alternate folding structure for the guide with positions 14-18 in the triplex loop base-paired with part of the pseudoknot, which would likely prevent RNP formation (Fig. S4A). Positively enriched mutations C15A and C17G were both predicted to disrupt this off-pathway structure (Fig. S4B). Comparing the predicted stability of the RNP-compatible ensemble of structures to the unconstrained ensemble shows that many triplex loop mutations are predicted to destabilize the unconstrained ensemble without disrupting the RNP-compatible ensemble (Fig. 1F). The triplex mutations are predicted to skew equilibrium toward the properly folded form and reduce the population of guide trapped in nonfunctional folded states, showing an alternate route to improved folding.

We also saw high tolerance to mutation in the extended stem, including positive enrichment of mutations expected to disrupt basepairing (Fig. 1B). To test whether extended stem hits could improve activity outside of the screen, we made a double mutant A88G and ^C63, which removes an asymmetric bulge while maintaining a single A:G noncanonical pair. We observed that the double mutation improved activity by >30% relative to Sv1 for both spacers (Fig. 1D).

We sought to identify alternatives to the U9C mutation to disrupt the polyU motif without reducing activity. We reverted the U9C mutation in Sv1 to generate Sv2, then mutated all three residues of the U11-U24:A95 triple in Sv2 to generate a C-G:C triple, resulting in Sv3. This modification increased activity for one spacer but decreased for another (Fig. 1G). Individual mutations at those sites in Sv1 essentially eliminated activity, which confirms the importance of triplex formation for guide function (Fig. 1D).

We added the beneficial single mutations C17G, A29C, ^G54 to Sv3, generating construct Sv5 that had greatly improved activity relative to Sv2 for both spacers (3.2-fold for spacer 1, 2.6-fold for Spacer 2, Fig. 1G). Including only mutations C17G, ^G54 and a truncated synthetic extended stem derived from the high-stability UvsX hairpin (Sv4) also showed significant improvement on Sv2, though lower activity than Sv5^25^. We tested two different versions of the extended stem with scaffold Sv5 – one that contained the A88G and ^C63 double mutant (Sv6), and one shortened version with the truncated synthetic UvsX hairpin (Sv7). Neither modified extended stem was significantly different from Sv5 in this assay (Fig. 1G). These results highlight scaffold Sv5 and its extended stem variants as a robust set of optimized sgRNA variants for CasX, achieving markedly enhanced editing efficiency and setting a foundation for subsequent protein engineering.

### Rational engineering and directed evolution of the CasX enzyme further improves potency

Having optimized the guide RNA, we next engineered the CasX protein. We first used an unbiased approach to identify beneficial amino acids. Plasmid recombineering was used to incorporate amino acid substitutions, deletions, and insertions into the WT PlmCasX sequence, generating a library predominantly composed of variants with a single mutation (Fig. S5)^26^.

Catalytically inactive versions of the library were generated by mutating each of the three conserved catalytic residues in the RuvC domain to alanine (D659A, E756A, or D922A), resulting in three libraries that could be used in repression-based screens^20^. This yielded a set of libraries that could be screened via different selection methods, allowing us to identify both consistently enriched mutations as well as those that are beneficial only in specific contexts.

The catalytically active version of the library was screened using a bacterial positive selection assay in which survival is coupled to cleavage of a target plasmid. Reasoning that PlmCasX might have low stability in human cells, given its origin from a bacterium identified in lower temperature groundwater samples, we ran the assay at 45 °C in addition to 37 °C to select for mutations that improve thermostability^19^. In parallel, the catalytically inactive libraries were screened for their ability to repress gene expression in a reporter *Escherichia coli* strain.

Mutations identified in multiple screens were selected for testing in human cells, along with a rationally designed mutation that introduced additional positive charge in the DNA-binding cleft of CasX. We measured their impact on editing of the *B2M* gene with multiple guide sequences when delivered as plasmid or as lentivirus under low multiplicity of infection (MOI) conditions. Three mutations, ΔP793, A708K, and L379R, were observed to improve activity guides via either plasmid or lentiviral delivery (Fig. 2A, S6). Combination of the three mutations resulted in a variant with greatly improved activity relative to the starting enzyme, referred to as Xv1 (Fig. 2A). Additional mutations recovered from screening were added to Xv1 to test for further improvements. Adding mutations F399L and I658V yielded a new variant, Xv2, with higher average editing than Xv1 and >10-fold activity improvement over PlmCasX with both guides (Fig. 2A).

**Figure 2.**
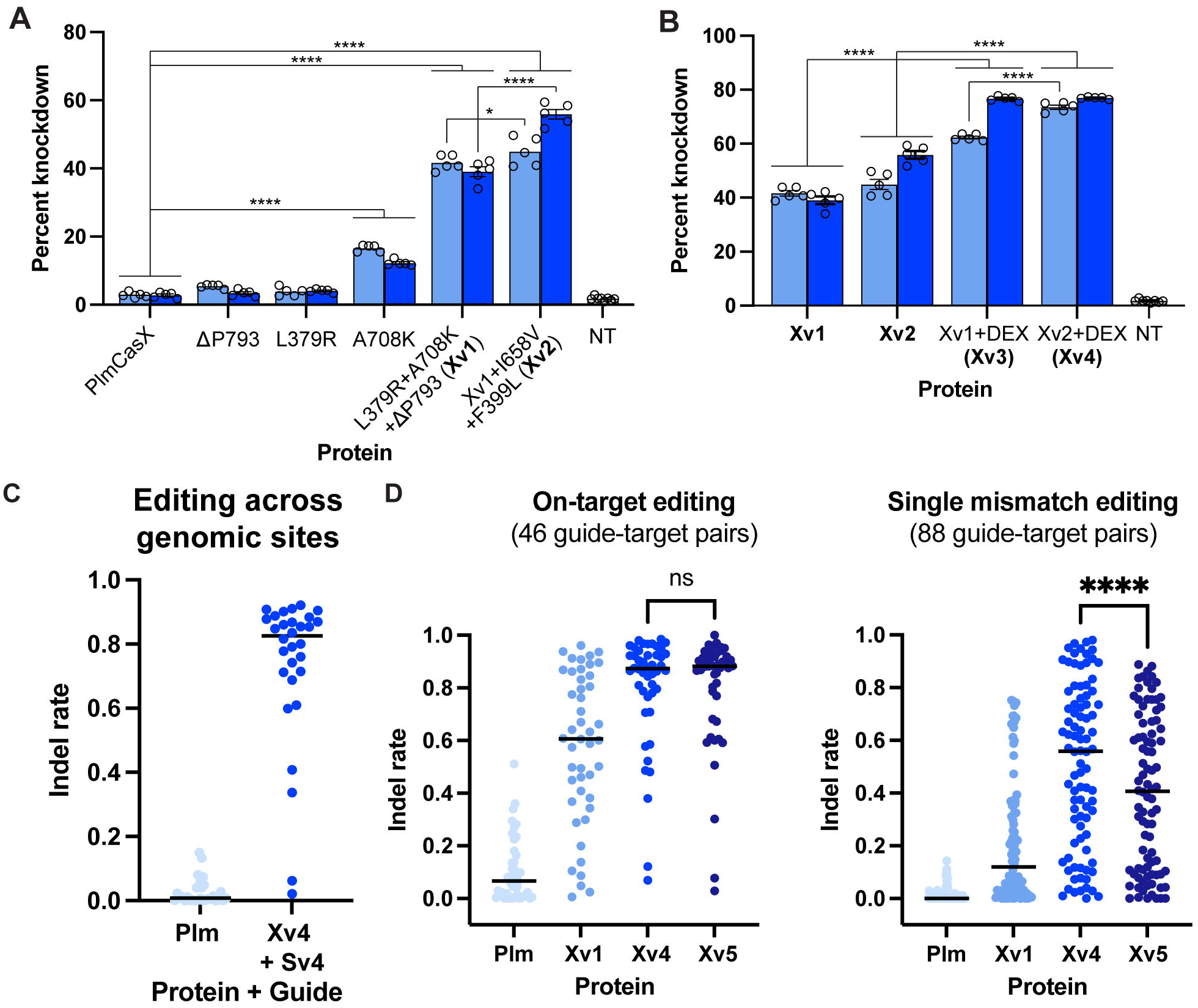
Engineering the CasX enzyme for improved editing. (A) HLA knockdown in human cells by WT and engineered CasX variants with two *B2M*-targeting guides via low-MOI lenti transduction. Three bioreplicates with three technical replicates each were performed. Mean, standard deviation, and individual data points for all replicates passing criteria is shown. NT = no treatment. p-values relative to the corresponding spacer for the indicated comparison are shown (one-way ANOVA, Dunnett’s multiple comparison test). * = p<0.05; **** = p<0.0001; ns = not significant (p>0.05). (B) HLA knockdown in human cells by PlmCasX-based engineered variants (Xv1 and Xv2) and those with Dpb NTSB and Helical IB domains (DEX) via low-MOI lenti transduction. Three bioreplicates with three technical replicates each were performed. Mean and standard deviation of all replicates passing criteria is shown. NT = no treatment. p-values are shown as in (A). (C) Average indel rate at 30 genomic sites, measured by NGS, of WT PlmCasX or Xv4 with scaffold Sv4. Each point represents the average editing across three replicates at a single genomic site. (D) Indel rates at on-target sites and single-mismatch off-target sites for the indicated CasX variant measured in pooled lentiviral screen. Each point represents the average editing across six replicates at a single spacer-target pair from the lentiviral library. ns = not statistically significant; **** = p<0.0001, paired t test.

Previous studies of PlmCasX and DpbCasX showed that the former more efficiently edits in human cells, but the latter cleaves DNA more efficiently *in vitro*.^20^ Based on this observation and the success other groups had exchanging small regions between CasX variants for improved activity,^22,23^ we explored exchanging full domains between WT DpbCasX and our engineered PlmCasX variants. Most chimeras between DpbCasX and Xv1 showed decreased activity when tested for editing in human cells, but the C-terminal region of the Helical I domain (Helical IB) and the complete Non-Target Strand Binding domain (NTSB) from DpbCasX maintained or improved activity when incorporated into Xv1 (Fig. S7). Exchanging both domains, which are contiguous in sequence, generated variant Xv3 with significant activity improvements over Xv1 (Fig. 2B). The same domain exchange was carried out for Xv2, which does not include any mutations in the altered domains. The resulting variant Xv4 shows improved activity over Xv2 for both spacers and over the Xv3 variant for one (Fig. 2B).

To assess the effects of both protein and guide engineering, Xv4 with an engineered scaffold was compared against WT PlmCasX with an unoptimized scaffold across 30 endogenous sites in human cells. The median indel rate increased from <1% with the minimally engineered enzyme and guide to 83% with engineered enzyme and guide, a >100-fold increase in editing at the average target site (Fig. 2C). We term the combination of highly engineered CasX enzyme and guide the X-Editor, or XE, system.

To determine whether the large increase in on-target editing that resulted from our engineering was accompanied by an increase in off-target editing, we designed a pooled screening system that enables the measurement of editing rates at dozens of on-target and single-mismatch off-target sites in parallel. As expected, the more active variants Xv1 and Xv4 show increased editing at mismatched targets as well as perfectly matched targets (Fig. 2D). We hypothesized that ΔP793 might have a disproportionate effect on off-target cleavage due to its location in the RuvC nuclease domain. The P793 residue was reintroduced, yielding variant Xv5. Xv5 reduced the mean editing rate at single-mismatch targets of 40%, as compared to 51% for Xv4, but shows no decrease in mean editing at matched targets (Fig. 2D). These results suggest that ΔP793 was a beneficial mutation in early rounds of engineering when overall activity was very low, but its benefits were outweighed by increased off-target effects in the context of a more active enzyme. Overall, the data demonstrates the XE RNP as a highly active editor with potential for therapeutic applications.

### Structural and biochemical characterization of XE platform

In order to understand the mechanisms by which our engineering efforts improved the activity of the XE RNP and inform future efforts, we performed cryo-EM of catalytically inactive Xv4-Sv7 RNP bound to a target DNA substrate and compared the resulting structure (global 2.77Å) to the structure of DNA-bound PlmCasX (Fig. 3A-B, Fig. S8)^22^. The two molecules overall showed high alignment (RMSD = 0.96Å) (Fig. S9A-B). Unlike the previous PlmCasX structure, the complex was captured in the absence of crosslinkers, and an extended DNA substrate enabled resolution of the complete R-loop, including reformed dsDNA at the PAM distal end and previously uncharacterized base stacking interactions (Fig. S9B, C)^22^.

**Figure 3.**
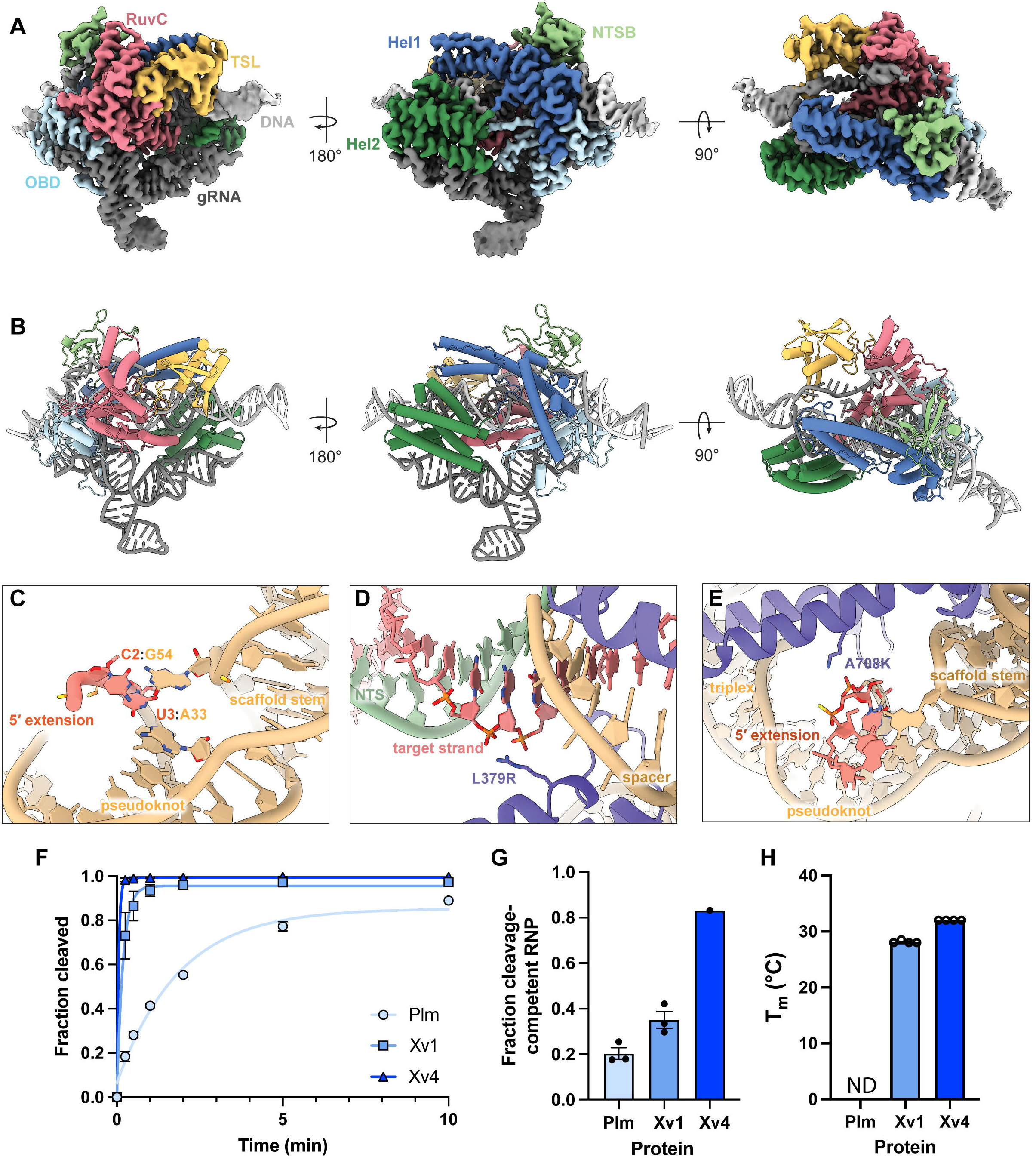
Mechanistic basis of improved potency of XE. (A-B) Cryo-EM density (A) and cartoon model (B) of XE (Xv4, Sv7) bound to target DNA. Colored by domain. (C) Novel base pairing interactions in the engineered gRNA scaffold. The 5′ extension is labeled in red, and new base pairs are indicated. (D) Interaction of L379R mutation with target strand of DNA. (E) Interaction of A708K mutation with 5′ extension of gRNA scaffold. (F) Kinetic cleavage assay performed with excess RNP over target DNA. Average substrate fraction cleaved and standard deviation at each time point is shown. The cleavage rate is modeled with a single exponential fit. (G) Fraction competence of indicated XE variant is shown, as determined by DNA cleavage with equimolar theoretical RNP and DNA substrate. Each point represents an independent experimental replicate. (H) Melting temperature of indicated apoenzymes as determined by fluorescent thermal shift assay. PlmCasX did not yield a clear fluorescence shift, potentially due to exposed hydrophobic regions even at low temperature.

Investigation of the guide RNA confirmed that U3 in the 5′ extension pairs with A33, as predicted based on enrichment and testing data (Fig. 3C). Unexpectedly, C2 pairs with G54, the nucleotide inserted into the bulge of the scaffold stem. ^G54 had the largest positive impact on activity of all the tested single mutations (Fig. 1D), which suggests that this novel interaction between the scaffold stem and the 5′ extension greatly benefits activity. This interaction may help pre-order the guide in a conformation amenable to RNP formation. The coordinated mutation of the second triple from U-A:U to C-G:C maintained the triplex as expected (Fig. S9D). The C17G mutation, along with the surrounding residues also found to be amenable to mutation, showed relatively weak cryo-EM density and no direct protein interaction, which is consistent with the hypothesis that the mutation functions by disrupting undesired folding states rather than through novel favorable interactions in the active RNP (Fig. S9E).

We next investigated the point mutations in Xv4 relative to PlmCasX. L379R forms a salt-bridge interaction with the phosphate backbone of the target strand immediately downstream of the R-loop (Fig. 3D). As one of relatively few protein-DNA interactions in this region, it might significantly improve DNA binding. A708K interacts with the phosphate backbone between nucleotides 3 and 4 of the guide RNA (Fig. 3E). Notably, this interaction would not be present in a version of the guide without a 5′ extension, which suggests that the benefit of the protein mutation is specific to the engineered guide. The mechanism by which ΔP793 affects activity and specificity is less clear. The deletion only modestly changes the local structure (Fig. S9F), but it may have a more significant impact at different stages of the target binding and cleavage process not captured in this structure, potentially by altering loop flexibility or repositioning K795 for non-target strand engagement in a manner that has differential effects on perfectly matched and partially mismatched targets.

Enzyme Xv1 was tested biochemically and compared to WT PlmCasX and found to cleave DNA more rapidly and form RNP more efficiently, consistent with improved DNA and gRNA binding from L379R and A708K, respectively (Fig. 3F-G). The apoenzyme also appeared to be more stable in a thermal shift assay, with a melting temperature of 28 °C while WT CasX was highly aggregation prone and failed to show a clear transition in the measured range (Fig. 3H, S10).

Xv4 was tested in the same assays and further improved stability, DNA cleavage, and RNP formation (Fig. 3F-H). The structure revealed that the chimeric NTSB and Helical 1B domains introduce several mutations at the interface with the target and non-target strands that might improve DNA binding (Fig. S9G-H). The observed stability enhancement of the apoenzyme may be driven either by domain exchange, the F399L and I658V point mutations, or both. Those mutations appear to induce only minor rearrangements of the local hydrophobic core but nonetheless improve activity despite a lack of clear mechanism (Fig. S9I-J). The surprising benefit of these mutations highlights the value of large-scale empirical screening.

These results confirm that our comprehensive engineering approach improved diverse properties important for effective gene editing. They also highlight the importance of an iterative approach to enable synergistic benefits, as seen in the 5′ guide extension that enabled interactions with new mutations in both the guide and protein that significantly improved activity. The structural data further serves as a basis for future engineering efforts.

### XE demonstrates efficient *in vivo* genome editing across multiple organ systems

To assess the versatility of the XE genome editing platform for *in vivo* applications, we evaluated its ability to edit a safe-harbor locus in mice. We first evaluated the capacity of XE to mediate editing in the retina. Adult mice were dosed via subretinal injection with an AAV8 vector expressing Xv4 under a modified version of a photoreceptor-specific promoter and guide Sv6 at multiple doses (Fig. 4A)^27,28^. Xv4 was used as Xv5 had not been developed at time of testing. At 2.5 weeks post-injection, indel analysis revealed editing of 66% in bulk retinal tissue at the highest dose, suggesting near complete editing of the targeted photoreceptors, which make up 70% of retinal cells (Fig. 4B)^29^.

**Figure 4.**
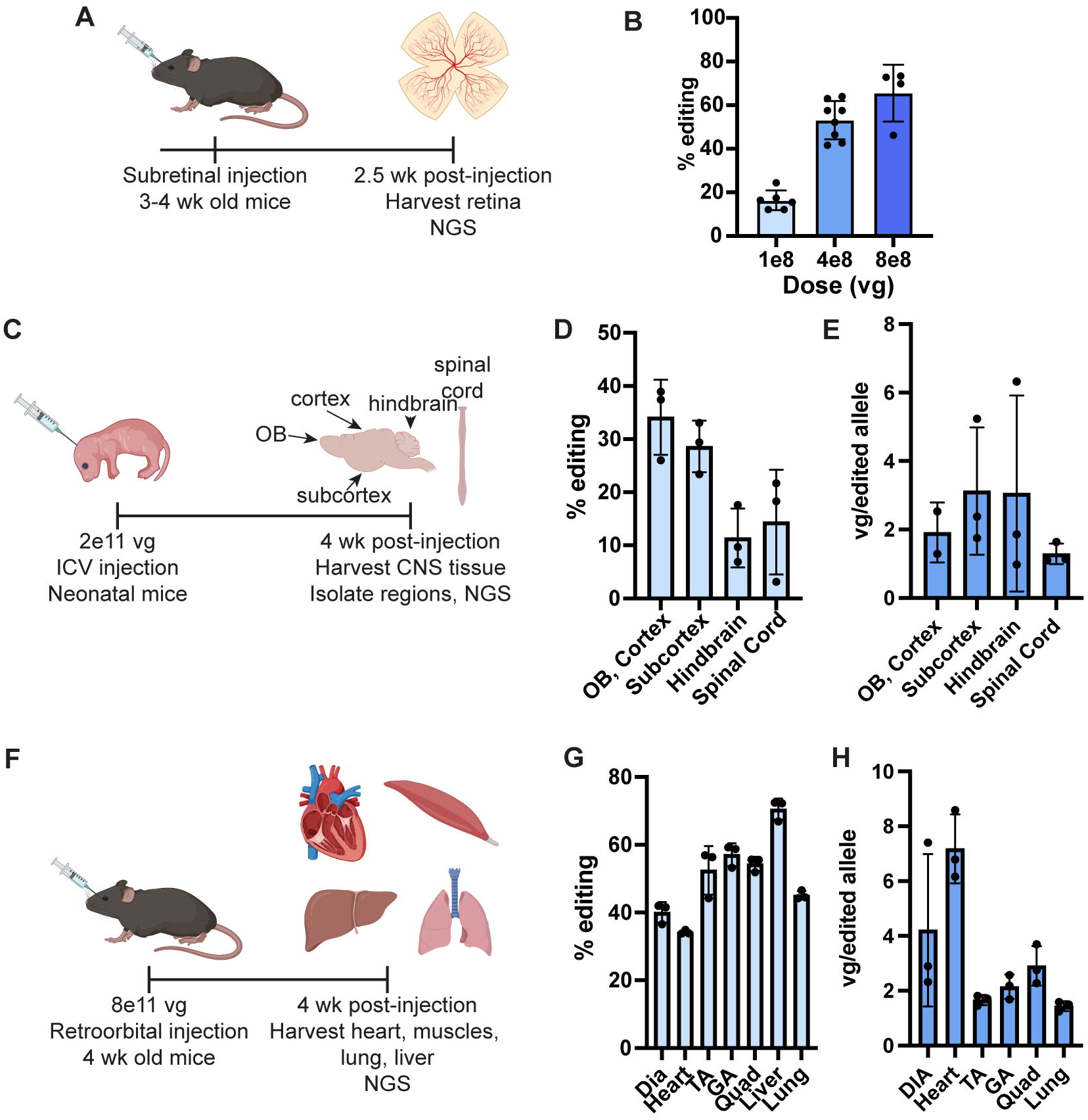
*In vivo* editing by local and systemic administration of AAV-XE. (A) Experimental design of retinal editing via local delivery in adult mice. (B) Editing rate at the ROSA locus in dissected retina as measured by NGS. Each point represents a retina. vg = viral genomes (C) Experimental design of central nervous system editing in neonatal mice. OB = olfactory bulb. (D) Editing rate at the ROSA locus in the indicated tissue type as measured by NGS. N=3 animals. (E) Estimate of viral genomes required for editing based on detectable AAV genomes in tissue after 4 weeks as measured by ddPCR. Calculated by dividing detected vg per haploid mouse genome by the fraction of edited alleles. (F) Experimental design for systemic editing in adult mice. (G) Editing rate at the ROSA locus in the indicated tissue. Dia = diaphragm. TA = tibialis anterior. GA = gastrocnemius. Quad = quadriceps. N=3 animals. Estimate of viral genomes required for editing, as in (E).

To evaluate editing in the CNS, we administered an AAV9 vector with Xv4 under a ubiquitous promoter to neonatal mice via intracerebroventricular (ICV) injection at a dose of 2.0e11 viral genomes (vg) (Fig. 4C). Four weeks post-injection, editing was assessed as well as the number of detectable viral genomes per diploid mouse genome (vg/dg), and the results were used to calculate the number of viral genomes per edited target allele (Fig. 4D, E). Across cortex, subcortex, hindbrain, and spinal cord, viral genomes per edited allele averaged from 1.3 to 3.1, indicating that few AAV episomes are required per cell for editing.

Finally, we tested whether XE could support efficient editing in muscle tissue following systemic administration. Adult mice received a single retroorbital dose of 8.0ell vg of AAV9 expressing Xv4 under a ubiquitous promoter, and editing outcomes were evaluated four weeks later (Fig. 4F). Editing rates were high across all tested tissues, reaching above 50% in all tested skeletal muscles, which compares favorably to other AAV-delivered CRISPR systems at higher doses (Fig. 4G)^30,31^. Due to the broad tropism of the AAV used, editing rates of 71% and 45% were seen in the liver and lung, respectively. Analysis of vg/edit showed averages ranging from 1.4 to 2.9 in the lung and skeletal muscle, with little editing increase in samples with higher vg/dg, suggesting saturation of targeted cell types (Fig. 4H, S11A). Editing in the heart closely aligned with the expected fraction of cardiomyocytes (30-40%), the cardiac cell type most effectively transduced by AAV9 (Fig. 4G)^32,33^. Higher editing rates were achieved via delivery to neonates via intravenous injection, with 55-70% editing observed across muscle tissues (Fig. S11B-C). Together, these studies show that the XE platform is highly active in every organ system tested.

### XE saturates editing of clinically relevant targets in non-human primates

We next performed a series of *in vitro* and *in vivo* proof-of-platform studies by evaluating Xv5 at targets relevant to cardiovascular disease. We chose to evaluate the editing potency of XE at the *PCSK9* and *APOC3* loci. Both genes are predominantly hepatocyte-expressed and are potential clinical targets involved in cholesterol and triglyceride homeostasis, respectively^34–37^. We first assessed XE editing activity at the *PCSK9* locus in primary human hepatocytes (PHHs) using LNPs formulated with Xv5 mRNA and a *PCSK9*-targeting gRNA. Sv7 was selected for LNP-based experiments, as its shorter length facilitates manufacturing. Dose-response studies doses were conducted, with editing rates measured by NGS (Fig. 5A). A Cas9-based base editor with a previously characterized *PCSK9*-targeting gRNA was used as a reference for potency^37^. XE demonstrated potent editing, with indel rates exceeding 80% (Fig. 5B). Furthermore, XE reached half-maximal editing at doses much lower than the base editor control. A similar study was carried out with *APOC3-*targeting gRNA, which also showed robust editing at low dose (Fig. S12).

**Figure 5.**
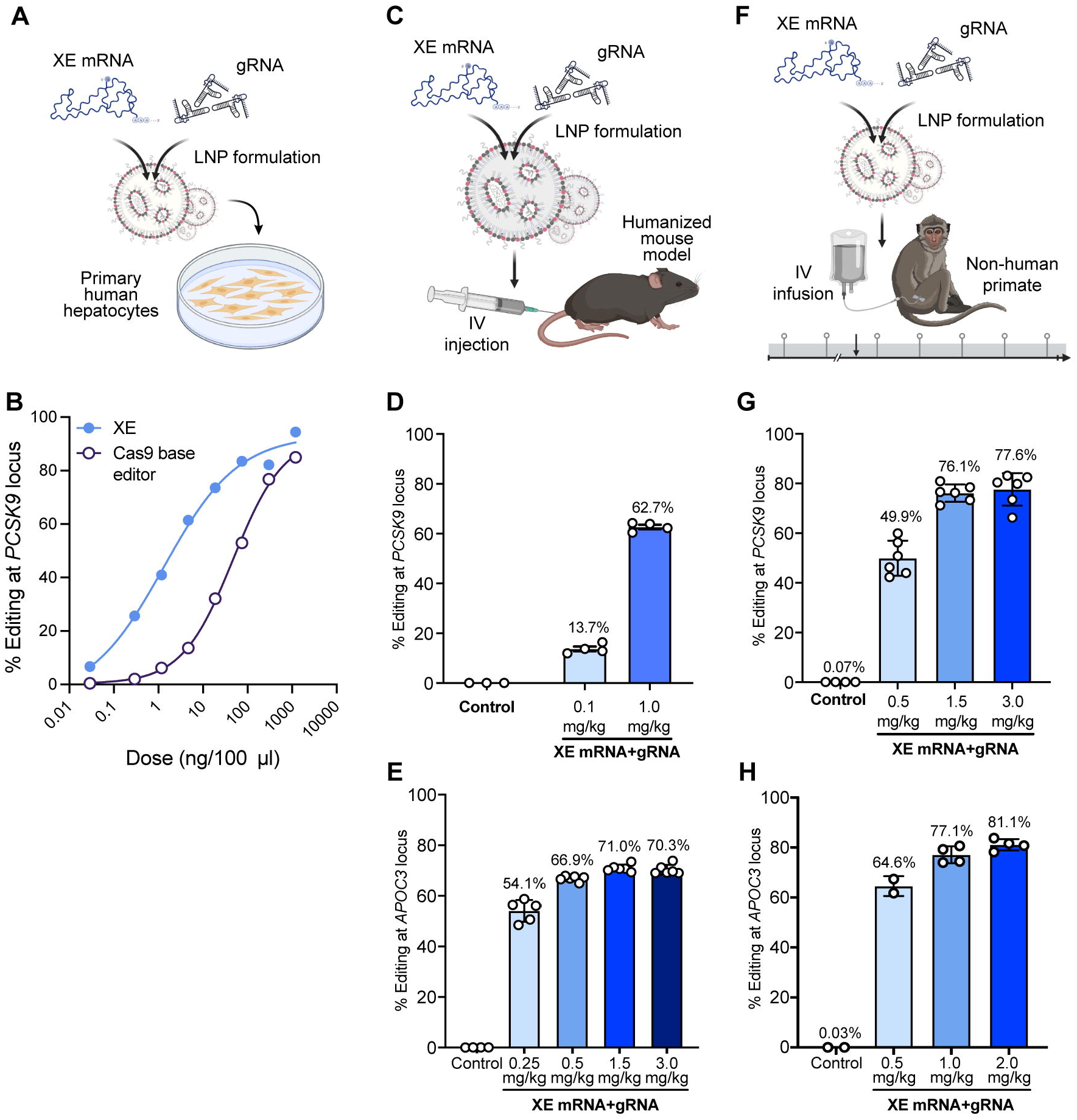
XE-mediated genome editing of *PCSK9* and *APOC3* in cynomolgus monkeys. (A) Schematic of primary human hepatocyte (PHH) editing by XE. (B) Editing of the *PCSK9* locus in PHH following treatment with LNP encapsulating either XE V5 mRNA and a *PCSK9*-targeting V7 gRNA or a Cas9 adenine base editor mRNA and a Cas9-compatible *PCSK9*-targeting guide. Editing was measured by NGS and dose response was modeled with a four-parameter fit. (C) Schematic of *in vivo* editing of *PCSK9* or *APOC3* in the respective mouse model, where either endogenous gene has been replaced by a human-derived sequence. (D) Editing rates at the *PCSK9* locus in humanized mice following injection of the indicated dose of LNPs encapsulating Xv5 mRNA and a *PCSK9*-targeting Sv7 gRNA. Control animals received phosphate-buffered saline (PBS). N=3 for PBS control, N=4 for each dose group. Individual points represent animals; bars show mean ± standard deviation. (E) Editing rates at the *APOC3* locus in humanized mice following injection of LNPs encapsulating Xv5 mRNA and *APOC3*-targeting Sv7 gRNA. N=4 for PBS control, N=5 for each dose group; bars show mean ± standard deviation. (F) Schematic of *in vivo* editing of *PCSK9* or *APOC3* in cynomolgus macaques. (G) Editing in liver tissue of cynomolgus macaques following a single intravenous infusion of *PCSK9*-targeting XE LNP. Control animals received PBS. Liver punch biopsies were collected at Day 60. N=4 for PBS control; N=6 for each dose group. Individual points represent animals; bars show mean ± standard deviation. (H) Editing in liver tissue of cynomolgus macaques following infusion of *APOC3*-targeting XE LNP. N=2 for 0.5 mg/kg dose, N=3 for 1.0 mg/kg, and N=4 for 2.0 mg/kg; bars show mean ± standard deviation.

To examine *in vivo* genome editing efficacy at the *PCSK9* locus, we employed a mouse model with a humanized gene. LNP was administered via intravenous injection, and sequencing of liver samples revealed 14% editing at 0.1 mg/kg and 63% at 1.0 mg/kg (Fig. 5C, D). Higher doses (1.5 and 3.0 mg/kg) showed editing efficiencies plateauing at ∼63-64%, consistent with the population of hepatocytes in murine liver, suggesting that saturation was reached at the 1.0 mg/kg dose (Fig. S13)^37^. The *APOC3*-targeting LNP was then assessed in a mouse model with a partially humanized *APOC3* gene. Following intravenous injection and sequencing of liver tissue, a 54% indel rate was identified at the lowest dose of 0.25 mg/kg, and editing rates exceeded 65% for doses of 0.5 mg/kg and above (Fig. 5E).

We next evaluated whether the same XE compositions could effectively edit in non-human primates. Cynomolgus monkeys received a single intravenous infusion of *PCSK9*-targeting XE LNP (Fig. 5F). Liver punch biopsies and sequencing analysis showed editing efficiencies averaging 50% at 0.5 mg/kg, 76% at 1.5 mg/kg, and 78% at 3.0 mg/kg (Fig. 5G). The similarity in editing between 1.5 and 3.0 mg/kg suggests saturation at or below the 1.5 mg/kg dose level, consistent with the expected population of hepatocytes in primate liver. Subsequent testing of lower doses showed editing of 68% at 0.75 mg/kg (Fig. S14).

The *APOC3*-targeting composition was assessed in cynomolgus macaques with a lower dosing regimen of 0.5, 1.0, and 2.0 mg/kg. Average editing efficiencies of 65%, 76%, and 81%, respectively, were observed, suggesting that XE edits nearly all hepatocytes at the 0.5-1 mg/kg doses (Fig. 5H). Collectively, these data establish XE as a potent genome editing system capable of high-efficiency editing across multiple targets, delivery modalities, tissues, and species.

### XE shows no off-target editing in primary human hepatocytes

To characterize the genome-wide specificity of XE with the two therapeutically relevant guides validated for high potency in vivo, we employed an off-target nomination strategy integrating three orthogonal approaches: in-cell detection (GUIDE-seq), *in vitro* biochemical profiling, and in silico prediction (Cas-OFFinder) (Fig. 6A)^38,39^. GUIDE-seq was performed in HEK293T cells with RNP electroporation^38^. For both guides, a single off-target site was identified with GUIDE-seq (Fig. 6B-C). An *in vitro* method based on cleavage of a library of synthetic potential off-target sequences nominated 283 sites for *PCSK9* and 172 for *APOC3* with enrichment values ≥1-fold over input library or ≥0.1% of on-target cleavage from a library of potential off-targets. Cas-OFFinder analysis identified 32 potential off-target sites for *PCSK9* and 147 for *APOC3* with four or fewer mismatches in the spacer or PAM sequence. Together, these three methods nominated roughly 300 candidate off-target sites for downstream validation for each guide (Fig. 6B-C).

**Figure 6.**
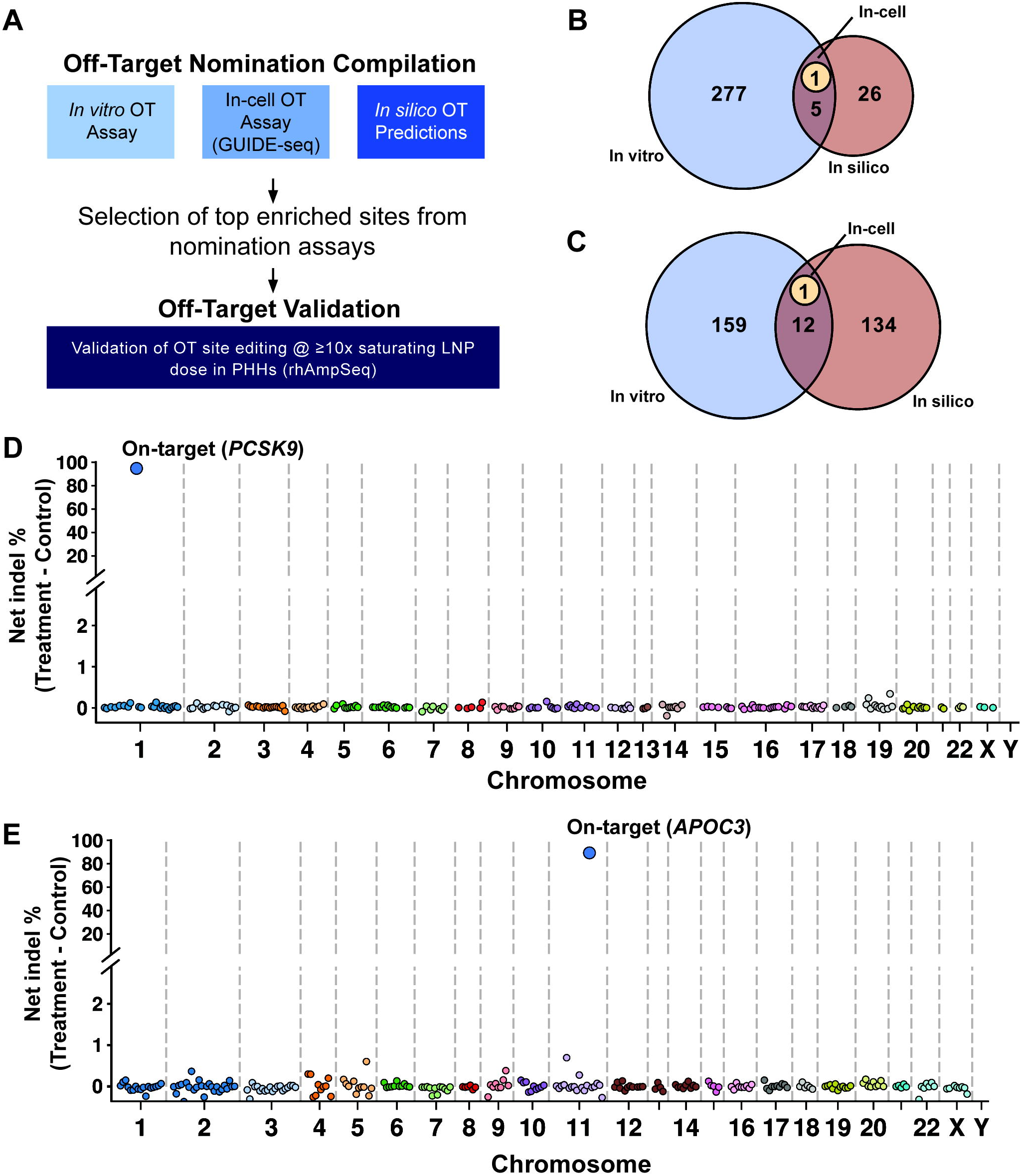
No off-target editing detected in primary human hepatocytes treated at supersaturating doses. (A) Schematic overview of the off-target nomination and validation strategy. (B-C) Venn diagram of sites identified by each of the off-target nomination strategies for *PCSK9* (B) and *APOC3* (C) guides. (D-E) Manhattan-style plots displaying indel rates at rhAmpSeq sites that passed quality control, shown for primary human hepatocytes treated with 10X EC90 dose of LNPs encapsulating Xv5 mRNA and *PCSK9*-targeting Sv7 gRNA (D) or 12X EC90 dose of LNPs with Xv5 mRNA and *APOC3*-targeting Sv7 gRNA (E). N=2 treated, N=1 untreated.

Validation of off-target editing was performed in PHHs treated with a supersaturating (≥10X EC90) dose of XE LNP. Pooled amplicon sequencing using rhAmpSeq was performed to evaluate the indel rate across all off-target sites for which amplicons could be designed. For any sites with editing above background, indel profiles were investigated to determine if they overlapped with the expected cut site at that target. Indels above background matching the XE editing profile were only observed at the *PCSK9* and *APOC3* on-target sites (Fig. 6D, 6E). For the *APOC3*-targeting guide, two sites had indel rates with statistical significance over untreated cells, but both showed indel profiles inconsistent with *bona fide* XE editing patterns, indicating they were likely due to sequencing error (Fig. S15). These findings demonstrate that XE maintains specificity even at supersaturating doses with guides capable of potent editing *in vivo*.

## DISCUSSION

Maximizing the impact of gene editing technologies requires that these technologies be exceptionally safe and effective. Generating a better therapeutic molecule means optimizing the CRISPR system for a fundamentally distinct set of requirements from those in which they evolved. In this study, we showcase systematic engineering of both the protein and gRNA components of the CasX system to overcome the inherent activity-specificity limitations of naturally occurring CRISPR enzymes.

Our systematic approach holistically interrogated and iteratively co-optimized both components of the CasX RNP enabling dramatic activity improvements while maintaining an exceptionally high specificity system. Between the protein and guide, we screened tens of thousands of variants in pooled screens and tested hundreds of designs in human cells. We identified meaningful improvements from rational changes, unbiased mutation screens, and large-scale domain exchanges. Our modifications led to improvements in RNP assembly, cleavage kinetics, and overall stability of both components, along with the destabilization of undesirable folded states of the guide. By combining holistic approaches as part of an iterative, results-oriented strategy, we improved many diverse functional characteristics in parallel, resulting in >100-fold activity improvements.

Recently, protein language and inverse folding models have allowed groups to sample evolutionarily or structurally plausible sequence space and generate novel functional enzyme variants. However, these approaches remain limited by their failure to incorporate data related to fitness for the activity of interest. Our approach complements these advances by demonstrating that comprehensive iterative empirical selection can optimize for both potency and specificity under conditions relevant to therapeutic genome editing, rather than relying on evolutionary sequence relationships alone. In the future, the combination of our large functional datasets with computational models may enable further improvements to the platform.

Our data demonstrate that not only is XE a dramatic improvement over naturally occurring CasX, it is a uniquely potent, specific, and flexible gene editor suitable for *in vivo* applications. XE demonstrates saturating editing of target cell types in multiple tissues using unoptimized AAV vectors and serotypes, generally requiring fewer than 4 viral genomes per edited allele. In NHP, XE reached saturating editing of the *PCSK9* locus at ≤1.5 mg/kg via LNP. This potency was exceeded at a second target, with saturating editing at the *APOC3* locus at a dose of 1 mg/kg. At doses of <1 mg/kg across two targets, XE shows significantly greater than was observed in comparable experiments for SpCas9-based compositions now in clinical trials, albeit with differences in target, formulation, and measurement approach (Table S1)^1,6,40^. This ability to saturate on-target editing at lower doses is particularly important to reduce exposure-dependent toxicities and inflammation associated with LNP delivery.^41^ Importantly, these potency gains were achieved without sacrificing genomic specificity. With no validated off targets even at doses far in excess of what is therapeutically necessary (≥10X EC90 in this study), XE now establishes a significantly broader therapeutic window for CRISPR approaches, enabling a wide range of safe and effective doses and increasing the likelihood that more challenging targets can be effectively addressed. This combination of enhanced potency and specificity is particularly relevant to preventive indications and common diseases, where the tolerance for treatment-related risks is substantially lower than in severe genetic disorders.

The XE platform also serves as a foundation for alternative CRISPR-based therapeutic modalities, including epigenetic silencing. A catalytically inactive version of the XE protein developed in this work serves as the DNA-targeting component of the Epigenetic Long-term X-Repressor (ELXR) platform, which enables targeted epigenetic repression without introducing DNA double-strand breaks. The ELXR technology has advanced into Phase 1 clinical development for the repression of *PCSK9* (ClinicalTrials.gov identifier: NCT07428473).^42^ Together, our findings establish that a comprehensive, iterative engineering approach can transform a naturally occurring CRISPR nuclease into a therapeutically optimized platform and provide a structurally characterized foundation for the development of increasingly potent and precise genetic medicines across diverse tissues and therapeutic applications.

## ACKNOWLEDGEMENTS

We thank David Savage, Jennifer Doudna, Jason Fernandes, and David Parrot for their critical review of the manuscript. We thank all members of Scribe Therapeutics for helpful discussions of this work. We thank Acuitas Therapeutics for providing the Lipid Nanoparticle (LNP) technology. We thank Katherine Baney, Han Spinner, Kian Taylor, Robert Teachenor, Raniel Alcantara-Lee, Lola Takhirov, Anna Mrak, Zuriah Meacham, and Angelica Castaneda for contributions to experiments.

## MATERIALS AND METHODS

### Structure annotations for guide scaffold base pairing and protein interactions

RNAPDBee was used to analyze pdb file 7WB1 to identify pairs that were base paired canonically or noncanonically using BPnet algorithm (accessed 11/22/2025), followed by manual annotation of the structure.^43^

Protein-interacting residues were identified with custom script to determine the distance between any atom in the base of each residue in the guide scaffold to any atom in the nuclease. Residues with a base within 3.5 A of any atom in the protein were identified as protein-interacting (Figure 1a).

### Guide library construction and screening

The designed guide scaffold library was ordered as a Twist oligo pool, PCR amplified, and cloned into lentiviral expression vectors including an expression cassette for Xv1 coupled to a puromycin resistance gene via a P2A sequence. A spacer sequence targeting GFP was then cloned into the guide construct. Lentivirus was produced in LentiX HEK cells according to standard protocols. Viral genome titer was determined by RT-qPCR. HEK 293T cells containing a GFP reporter were transduced with lentivirus at <0.5 multiplicity of infection (MOI). After 48 hours, puromycin was added and cells were selected for 4-6 days. Cells were allowed to recover in nonselective media before being trypsinized, resuspended in PBS with 4′,6-diamidino-2-phenylindole (DAPI), and subjected to fluorescence-activated cell sorting (FACS) for the isolation of GFP– cells. A sample of unsorted cells was also collected to establish a naïve baseline. Genomic DNA was isolated and scaffolds were sequenced via next-generation sequencing (NGS). Scaffold sequences were aligned to the Sv1 reference sequence to determine identity. Read counts for each scaffold were normalized for sequencing depth and enrichment of normalized abundance relative to a naïve sample was calculated. The full screen, from lentiviral production through genomic isolation and screening, was carried out in duplicate. For each variant, a weighted average across the two screens was calculated with weighting based on relative representation in the naïve population for each screen. Ensemble folding energies and minimum free energy structures were predicted using RNAfold (http://rna.tbi.univie.ac.at/cgi-bin/RNAWebSuite/RNAfold.cgi).

### Protein library construction and screening

Libraries of CasX protein variants were constructed using plasmid recombineering as previously described^26^. In addition to a library built on the catalytically active PlmCasX, three libraries were constructed on base molecules each having an inactivating mutation to one of the residues of the RuvC catalytic triad. A fourth library was constructed with all three inactivating mutations in combination.

Screening of the catalytically inactive libraries was carried out in an *E. coli* strain carrying monomeric Red Fluorescent Protein (mRFP) and Superfolder Green Fluorescent Protein (sfGFP)^44^. Cells were transformed with plasmid expressing the CasX library under anhydrotetracycline-inducible promoter and guide RNA with either a GFP-targeting or non-targeting spacer. Transformed cells were then either plated on LB agar with appropriate antibiotic and aTc and RFP–/GFP+ colonies identified visually and picked or grown in liquid culture and sorted for RFP–/GFP+ cells via FACS. Plasmids were isolated and sequenced to identify selected variants. Screening of the active library was performed using the CcdB selection assay as previously described.^45^ Mutations identified in these screens were selected for further characterization.

#### Lipofection of guide and protein variants

Scaffold and protein variants were cloned into plasmids expressing a guide RNA driven by a U6 promoter and an XE variant coupled to mScarlet by a P2A tag, expressed by a UbC promoter. All scaffold variants were tested with WT PlmCasX, and all protein variants were tested with scaffold Sv6. All variants were tested with two *B2M*-targeting guides, and a subset were also tested with two additional lower activity guides, also targeting *B2M*.

HEK293T cells were seeded in 96-well plates at a density of 10,000 cells/well and transfected with 100 ng plasmid using Lipofectamine 3000 (Thermo Fisher Scientific, cat. no. L3000015) according to the manufacturer’s protocol followed by centrifugation at 1000g for 10 minutes. At 24h post transfection, a media exchange was performed using 1X DMEM + GlutaMax (Gibco, cat. no. 10565-011) supplemented with 10% (v/v) heat-inactivated bovine fetal calf serum (Corning, cat. no. 35-016-CV) and 100 U ml-1 Penicillin-Streptomycin (Gibco, cat. no. 15140-122). On day 7 post transfection, cells were stained with 0.03ug Alexa Fluor 488 anti-human HLA-A,B,C Antibody (BioLegend, cat. no. 311413) in 50uL staining buffer (1X DPBS (VWR Life Science, cat. no. 02-0119-0500) and 3% heat-inactivated bovine fetal calf serum (Corning, cat. no. 35-016-CV). The cells were washed in 1X DPBS and stained with DAPI (Invitrogen, cat. no. 62248).

Cells were analyzed by flow and gated for mScarlet positive cells, which was used as a transfection control. Only mScar+ cells were analyzed for HLA depletion. Only replicates where the percentage of mScar+ cells was greater than 40% and the total number of measured mScar+ cells was greater than 5000 were analyzed to exclude inefficient transfections. All analyzed variants were measured in at least two biological replicates with two or more technical replicates meeting inclusion criteria. To assess significance, one-way ANOVA with Dunnett’s multiple comparison test were performed. Variants that differed from the reference molecule with a p-value < 0.05 were considered significant.

#### Lentiviral transduction of protein variants

Lentivirus was produced by co-transfecting Lenti-X 293T cells with transfer plasmids and packaging vectors (psPAX2, pMD2.G) using TransIT-LT1 (Mirus, MIR 2300). Viral supernatants were concentrated with 36% PEG6000 in 1X DPBS. Functional titer was determined by performing transductions across a dilution series and measuring mScarlet + cells.

HEK293T cells were seeded in 96-well plates at a density of 10,000 cells/well. After 24h, cells were transduced with the amount of lentivirus calculated to achieve a MOI of 0.5 based on functional titer, followed by centrifugation at 1000g for 10 minutes. Cells were passaged on day 4 and harvested and stained on day 7 in the manner described above.

Cells were analyzed by flow as described above, except that only replicates with less than 40% mScar+ cells were considered to limit the likelihood that a substantial number of cells have more than one integrant.

#### Comparison of PlmCasX to XE on endogenous targets

Thirty spacers were designed targeting six human genes, with five spacers per gene. Plasmids were generated either with PlmCasX with a version of the guide containing a 4-nt 5’ extension relative to the published sequence (Sv0) or with protein Xv4 with Sv4, with a puromycin resistance gene coupled to the CasX variant by a P2A tag. 100ng of plasmid was transfected into ∼60K adherent HEK293T cells in flat bottom 96 well plates in duplicate using Lipofectamine 3000. 1.5ug/ml puromycin was added 24 hours after transfection and selection occurred over 4 days. On the fifth day, the duplicates were pooled and the genomic DNA was harvested using Zymo Quick-DNA 96 kit. The gDNA from each round was amplified for Illumina sequencing and indels were identified using the CRISPResso2.0 package^46^. Three experimental replicates were carried out and average editing at each site across replicates was determined.

#### Pooled activity and specificity assay

A pooled library of lentiviral vector plasmids was generated of on- and off-target spacer-target pairs with multiple PAM sequences. Only the sites with TTCN PAMs are analyzed in this study, representing 48 on-target sites and 88 single-mismatch off-target sites. Each plasmid contained a GFP expression cassette, an Sv4 guide sequence with a U6 promoter, an associated target site, and a unique identifier sequence. The plasmid was used to generate a lentiviral library according to standard methods. HEK293T cells were transduced at low MOI and transduced cells were isolated by flow cytometry. Cells were adapted to suspension culture by serum reduction.

Cells were seeded in 96-well deep well format and transfected with 500 ng of plasmid expressing engineered XE proteins and a puromycin resistance gene using PEI Max (Kyfora Bio). 2 μg/mL puromycin was added at day three post-transfection. At day eight, treated cells were collected and lysed, and genomic DNA was extracted using a genomic DNA isolation kit (Zymo Research). Genomic DNA was then PCR amplified with custom primers to generate amplicons compatible with Illumina NGS and sequenced on a NextSeq instrument. Sample reads were demultiplexed and filtered for quality. The fraction of reads with indels was then quantified for each spacer-target. For each sample and spacer-target pair, data based on <500 reads were removed. Fraction indel values for each sample and spacer-target pair were subtracted by the average fraction indel value across Cas9-treated samples with the same spacer-target pair, where Cas9 served as a negative control due to incompatibility with the XE guide RNA.

#### Protein Expression and Purification

The expression plasmid encoding the variant CasX protein coding sequence was transformed into the *E. coli* BL21* strain. Cells were grown in Terrific Broth (TB) at 37°C to an OD600 of 0.5, and culture was cooled at 4°C until OD600 reached ∼2 and the culture temperature was 16°C at which point 1mM IPTG was added. Cells were grown at 16°C for 16-24 hours and harvested by centrifugation. Bacterial pellets were resuspended in lysis buffer (50 mM HEPES-NaOH, 250 mM NaCl, 5 mM MgCl2, 0.5 mM TCEP, 10% Glycerol, pH 8, 1mM Benzamidine-HCL, 1 mg/mL Lysozyme, 0.1% CHAPS) and 1mM PMSF was added prior to lysis by homogenization. The cell lysate was clarified by centrifugation and the supernatant was applied to a Heparin 6 FF column pre-equilibrated with Buffer A (50 mM HEPES-NaOH, 250 mM NaCl, 5 mM MgCl2, 0.5 mM TCEP, 10% Glycerol, pH 8). The column was washed with 5 column volumes (CV) of Buffer A, followed by 3 CV of Buffer B (50 mM HEPES-NaOH, 500 mM NaCl, 5 mM MgCl2, 0.5 mM TCEP, 10% Glycerol, pH 8), and sample was eluted with 1.75 CV of Buffer C (50 mM HEPES-NaOH, 1 M NaCl, 5 mM MgCl2, 0.5 mM TCEP, 10% Glycerol, pH 8). The sample was then applied to a StrepTactin HP column pre-equilibrated with 2CV of Buffer B. The column was washed with 10 CV of Buffer B and eluted with 1.65 CV Buffer D (50 mM HEPES-NaOH, 500 mM NaCl, 5 mM MgCl2, 0.5 mM TCEP, 10% Glycerol, pH 8, 2.5 mM Desthiobiotin).

Finally, the eluate was applied to a Superdex 200 pg column pre-equilibrated with Buffer E (50 mM HEPES-NaOH, 250 mM NaCl, 250 mM KCl, 0.5 mM TCEP, 10% Glycerol, pH 8). An isocratic elution was performed with Buffer E. CasX fractions were pooled and protein concentration was measured by a Pierce 660 nm Protein Assay. TEV was added at 1:100 ratio to CasX along with 1 mM DTT and incubated overnight at 4°C. The TEV cleavage reaction was concentrated using 50kDa Amicon spin concentrators and further purified using size exclusion chromatography on a HiLoad 26/600 Superdex 200 pg column pre-equilibrated with Buffer F (25 mM NaPO4, 300 mM NaCl, 1 mM TCEP, 10% Glycerol, pH 7.25). An isocratic elution was performed with Buffer F, CasX fractions were pooled and concentrated using a 50 kDa Amicon spin concentrator. Protein concentration was measured using a Pierce 660nm assay.

#### RNP complex assembly and purification

RNP complexes were assembled using recombinant CasX apoenzyme and sgRNA produced by solid phase oligonucleotide synthesis (Trilink). Lyophilized sgRNA was reconstituted in 1 mM KPO4, pH 6.5 buffer at 100 µM before being folded by heating to 80°C for 15 minutes and cooling to room temperature on the bench for 30 minutes. The RNP complexation was assembled on ice in buffer containing 25 mM KPO4, 300 mM NaCl, 1 mM MgCl2, and 10% Glycerol , pH 7.5, with 20 µM apoenzyme and 24 µM sgRNA. The complexation reaction was incubated on ice for 15 minutes, then 10 minutes in 25, 30, and 37°C water baths, sequentially, before being returned to ice. The reaction was diluted 1:1 with Dilution Buffer (25 mM KPO4, 10% Glycerol) and purified by anion exchange chromatography using a Capto HiRes Q 10/100 column (Cytiva) pre-equilibrated with MPA (25 mM KPO4, 150 mM NaCl, 10% Glycerol, pH 7.5). The column was first washed with 5 CV of 25% MPB (25 mM KPO4, 1 M NaCl, 10% Glycerol, pH 7.5), and then RNP complexes were eluted with a two-phase linear gradient, 25-45% MPB over 10 CV, followed by 45-65% MPB over 5 CV. Finally, any remaining material was eluted with 100% MPB for 5 CV. Monomeric RNP complexes were pooled and concentrated by ultrafiltration using 15 mL, 50 kDa MWCO Amicon spin concentrators (Millipore), was injected onto a HiLoad 16/600 Superdex 200 prep grade column (Cytiva) equilibrated with MPC (25 mM KPO4, 300 mM NaCl, 0.5 mM TCEP, 10% Glycerol, pH 7.5). The monomeric RNP peak was pooled on ice and concentrated by ultrafiltration again. The RNP was concentrated to 40 µM, as assayed by Pierce 660 nm protein assay (ThermoFisher), before being diluted 1:1 with dilution buffer, aliquoted, snap frozen in liquid nitrogen, and stored at -80°C.

#### Substrate-engaged complex assembly and purification

DNA substrates were ordered as ssDNA oligos (IDT) and resuspended at 100 µM in 10 mM Tris-HCl, pH 8. Complementary strands were annealed by mixing 1:1 with a final buffer composition of: 50 µM each ssDNA, 10 mM Tris-HCl, 50 mM NaCl, pH 8. Samples were heated to 95°C in a thermalcycler for 5 minutes before being allowed to cool to room temperature on the benchtop. Purified RNP complexes were thawed on ice before being mixed with substrate at a 1:1.5 (RNP:substrate) ratio in binding buffer (25 mM KPO4, 150 mM NaCl, 5 mM MgCl2, 10% glycerol, pH 7.5). Reactions were incubated in a 25 °C water bath for 30 minutes, then cooled to 4°C. Assembled complexes were purified by SEC using the same method as the RNP complexes alone. The peak containing substrate-engaged RNP was pooled and concentrated by ultrafiltration, ending at 50 µM. This was diluted in half with nuclease-free water to yield 25 µM substrate-engaged RNP complexes in Storage Buffer (12.5 mM KPO4, 150 mM NaCl, 0.25 mM TCEP, 5% Glycerol, pH 7.5).

#### Cryo-EM sample preparation and data collection

Cryo-EM samples were prepared with the target-bound RNP (7 µM final concentration). Samples (4 µL) were applied to Quantifoil R2/1, 300 mesh, holey carbon copper grids (Electron Microscopy Sciences) after glow discharging the grids at 11 mA for 40 sec (Pelco). CHAPSO detergent was added to a final concentration of 0.15% just before grid freezing to ensure uniform ice thickness and particle distribution. Samples were vitrified with a Vitrobot Mark IV system (Thermo Fisher Scientific) maintained at 10°C and 100% humidity. Grids were blotted at blot force 1 for 7 sec and plunge frozen in liquid ethane.

Grids were screened for optimal ice thickness and particle concentration. Data collection was carried out from a single screened grid using a 300 kV Titan Krios cryo-transmission electron microscope (Thermo Fisher Scientific) equipped with a K3 camera (Gatan) and an imaging energy filter (Gatan) operated at a slit width of 20 eV. The dataset was collected in counting super-resolution mode with a nominal magnification of 105,000x leading to a physical pixel size of 0.832 Å (super-resolution pixel size is 0.416 Å). The data were collected at a dose rate of 24 e-/pixel/sec with a total electron dose of 48 e-/Å2 applied over 40 frames and a targeted defocus range of -0.8 µm to -1.8 µm.

#### Cryo-EM data processing and model building

The cryo-EM data processing workflow was carried out in CryoSPARC v5.^47^ Movie frames were motion corrected using Patch motion correction and binned two-fold to yield a motion-corrected micrograph stack with a pixel size of 0.832 Å. Micrographs were manually curated and an initial round of particle picking was performed using blob picker on a subset of 1,000 micrographs. A 2D classification job on the initial particle stack yielded 2D templates that were used for template-based particle picking on the entire dataset. The template-picked particle stack was curated using 2D classification and used for *ab initio* reconstruction and heterogenous refinement. Further analysis of the particle stack was done by iterative 3D classification with a focus mask and non-uniform refinement. The final particle stack was polished using Reference-based motion correction and used for local refinement to create the final reconstruction. Final reconstruction was uploaded to the Electron Microscopy Data Bank (EMDB).

Model building was done in Coot.^48^ For the CasXE protein, a starting model was generated with AlphaFold3,^49^ and rough domain fitting was performed in ISOLDE^50^ using a gaussian blurred map. For the sgRNA and DNA substrate, nucleic acid chains from 7WAY were used as starting models and mutated in Coot. After initial rigid-body docking, residues were manually fit into the high-resolution map using real-space refinement modules. Rounds of real-space refinement in PHENIX^51^ were carried out and the outliers were manually corrected in Coot before deposition in the Protein Data Bank (PDB). Structures were visualized and figures generated with ChimeraX^52,53^.

#### Biochemical cleavage assays

Target probes containing the B2M spacer 2 sequence were generated by annealing 60-nt DNA oligonucleotides with 5′ fluorescent labels (IRDye700 and IRDye800 on the non-target strand and target strand, respectively). Oligonucleotides were mixed in a 1:1 ratio in 1x cleavage buffer (20 mM Tris HCl pH 7.5, 150 mM NaCl, 1 mM TCEP, 5% glycerol, 10 mM MgCl2), heated to 95° C for 10 minutes, and allowed to cool to room temperature. XE RNPs were formed with the indicated XE and Sv4 guides at a final concentration of 1 µM with 1.5-fold molar excess guide in 1x cleavage buffer at 37° C for 10 min before being moved to ice.

For kinetic cleavage assays, cleavage reactions were run at 37° C with 200 nM RNP complex and 10 nM target probe. Time points were taken at 0.25, 0.5, 1, 2, 5, and 10 minutes, and quenched by the addition of an equal volume of quench buffer (95% formamide, 20 mM EDTA pH 8.0). Samples were denatured at 95° C for 10 minutes and then analyzed via denaturing polyacrylamide gel electrophoresis (PAGE) with 7 M urea and 10% acrylamide. Gels were imaged on a Typhoon gel scanner and analyzed with ImageQuant (Cytiva). Fraction cleaved of the non-target strand was determined via the fraction of signal per lane present in degradation products vs full-length substrate. Cleavage rates were determined in Prism using a one-phase association fit through all replicate data.

For determination of fraction cleavage-competent RNP, cleavage reactions were carried out at 37°C with 100 nM of both RNP and target probe. Time points were taken at 5, 10, 30, 60, and 120 minutes. Samples were quenched, subjected to PAGE, and imaged as above. The amount of cleaved target was observed to plateau over this timescale, confirming that CasX and XE act essentially as a single-turnover enzyme under the assayed conditions. Cleavage traces were fit in Prism with a biphasic rate model, as a monophasic model provided a poor fit under this concentration regime. The calculated plateau, expressed as a fraction of target cleaved, was used as the estimated fraction of cleavage-competent RNP, assuming that each active RNP molecule bound and cleaved a single target molecule. The plateau was calculated for each replicate independently, then plateaus were averaged.

#### Thermal shift assay

Thermal shift assays were carried out by adding CasX apoprotein to a final concentration of 2 μM in thermal shift buffer (20 mM HEPES pH 7.5, 400 mM NaCl, 1 mM TCEP, and 0.1% β-D-octyl glucoside) kept on ice. Samples were prepared in quadruplicate. GloMelt dye (Biotium) was added according to manufacturer recommendations. Samples were transferred to a BioRad qPCR machine. Fluorescence of the dye was measured using the SYBRGreen channel starting at a temperature of 5 °C; temperature was increased at a rate of 1 °C/min to a final temperature of 75 °C. For each replicate, the change in fluorescence intensity with respect to temperature was determined (dRFU/dT), and the maximum of the derivative was used to determine melt temperature. A Tm was not determined for WT PlmCasX as it showed high fluorescence at low temperature and did not display a clear transition as was seen for the engineered variants.

#### AAV mouse experiments

AAV8 vectors for retinal editing coding for Xv4 under a truncated GRK1 promoter and a ROSA-targeting Sv6 guide were produced and purified by affinity chromatography. Vectors were titered using ddPCR. 3-4 week old C57BL6J mice were injected subretinally with 1.0e8, 5.0e8, or 8.0e8 vg of AAV. 2.5 weeks following injections, mice were euthanized and retinae were dissected and harvested for gDNA. Samples were subjected to NGS analysis of indels at the target site.

AAV9 vectors for CNS editing coding for Xv4 under a U1a promoter and a ROSA-targeting Sv6 guide were produced and purified by affinity chromatography. Vectors were titered using ddPCR. P0 neonate C57BL6J mice were given an intracerebroventricular injection of 2e11 vg of AAV. 4 weeks following injections, mice were euthanized, brains were harvested and dissected. gDNA was harvested. NGS analysis of indels at the target site was performed. Viral genome per diploid genome (vg/dg) was determined by performing ddPCR with probes specific to Xv4 (to quantify viral genomes) and RPP30 (to quantify mouse genomes). Viral genomes per edited allele were determined by dividing vg/dg value by 2 and then dividing by the indel rate for that sample determined by NGS.

AAV9 vectors for muscle editing coding for Xv4 under a UbC promoter and a ROSA-targeting Sv6 guide were produced and purified by affinity chromatography. Vectors were titered using ddPCR. For the neonatal experiment, 1e12 vg were delivered by facial vein injection to P0-P1 C57BL6J mice. Tissues were harvested four weeks after injection and indels at the target locus were quantified by NGS. 4-5 week old mice were dosed with 8e11 vg via retroorbital injection and tissues were harvested 4 weeks post injection. Indel analysis, vg/dg, and vg/edited allele were determined as described above.

#### LNP formulation

mRNA and gRNA were formulated into lipid nanoparticles (LNPs). For select studies, LNPswere developed and formulated by Acuitas Therapeutics (Vancouver, Canada).

#### Primary Human Hepatocyte (PHH) culture and screening

Cryopreserved Primary Human Hepatocytes (PHH; Lonza, lots HUM190131 and HUM181441) were thawed following the manufacturer’s instructions and plated at 60,000 cells per well in collagen I–coated 96-well plates (Corning® BioCoat®). After 24 h, the medium was replaced with PHH Maintenance Medium prior to lipofection. PCSK9- and APOC3-targeting gRNAs were mixed with XE V5 mRNA at a 1:1 mass ratio and formulated into LNPs. LNPs were administered across a 4-fold dilution series, with 1200 ng total RNA as the top dose.

Genomic DNA was extracted from 100 µL of DNA/RNA Shield lysate using the Quick-DNA™ 96 Plus Kit (Zymo Research) and quantified with the Qubit™ dsDNA HS Assay Kit (Thermo Fisher Scientific). PCR1 amplification of PCSK9 target loci was performed with KAPA HiFi HotStart ReadyMix (Roche) using locus-specific primers. PCR1 products were re-amplified in PCR2, and libraries were sequenced on an Illumina platform. Indel frequencies were quantified using CRISPResso2 (Clement et al., 2019, Nat Biotechnol 37:224–226).

#### LNP mouse experiments

In vivo studies were conducted according to IACUC guidelines using commercially available humanized transgenic PCSK9 mouse model (B-hPCSK9-Plus, human *PCSK9* transgenic mouse, Biocytogen, Catalog number 112751; Strain name: C57BL/6-*Pcsk9^tm2(PCSK9)Bcgen^*/Bcgen). In this model, the genomic region of the mouse *Pcsk9* gene that encodes the full-length PCSK9 protein, as well as the 5’UTR and 3’UTR, are replaced by human *PCSK9* gene sequence and 5’UTR and 3’UTR sequences. LNP formulations encapsulating Cas-XE mRNA and *PCSK9* targeting sgRNA were delivered via a single tail vein intravenous bolus injection to 10 to 12 week-old B-hPCSK9 mice. Mice were divided into groups and dosed with vehicle or LNPs at low doses of 0.1 and 1.0 mg/kg or higher doses of 1.5 and 3.0 mg/kg. At seven days post injection, mice were sacrificed and their livers collected for on-target *PCSK9* gene editing analysis via NGS and CRISPResso analysis.

To assess editing at the human *APOC3* locus in vivo, eight-week-old B-hAPOC3 humanized mice (C57BL/6N-*Apoc3^tm1(APOC3)Bcgen^*/Bcgen) were administered *APOC3*-targeted LNPs at 0.25, 0.50, 1.5 and 3.0 mg/kg dose levels via a single tail vein intravenous bolus injection. In this model, exons 2-4 of the mouse *APOC3* gene that encodes full-length protein are replaced by human *APOC3* exons 2-4. In this study, liver tissue was harvested 14 days post dose for on-target *APOC3* gene editing.

#### NHP experiment

A study in cynomolgus monkeys was performed at Altasciences according to IACUC guidelines. Male and female monkeys (2-5 years old at initiation of dosing) were genotyped for spacer sequence identity prior to randomization and then were randomized to study groups according to genotyping results and pre-established social units. Monkeys were administered premedication (diphenhydramine, 5 mg/kg; dexamethasone, 1 mg/kg; famotidine 0.5 mg/kg) approximately 24 hours and 1-2 hours prior to dosing with vehicle or a LNP formulation encapsulating XE mRNA and sgRNA targeting either the *PCSK9* or the *APOC3* cyno locus. For *PCSK9* targeting, monkeys were divided into three groups for LNP administration at 0.5, 1.5, and 3.0 mg/kg dose levels. A subsequent experiment was carried out at 0.5 and 0.75 mg/kg doses. For *APOC3* targeting, monkeys were divided into three groups for LNP administration at 0.5, 1.0 and 2.0 mg/kg dose levels. LNP formulations were administered via intravenous infusion over 1 hour.

An ultrasound-guided needle punch biopsy was performed on Day 84 post-dose for the full dose-range *PCSK9*-targeting experiment, Day 15 for the lower dose *PCSK9*-targeting experiment, and Day 18 for the *APOC3*-targeting experiment. The liver biopsy sample was flash frozen and shipped to Scribe Therapeutics for DNA sequencing analysis. The target site was PCR-amplified, sequenced via NGS, and analyzed for *APOC3* editing using CRISPResso.

#### GUIDE-seq

In preparation for nucleofection, CasX apoenzyme and 2-fold molar excess guide RNA were combined and complexed by incubation at 37°C for 15 minutes. 100 pmol of complexed RNP was then combined with 200 pmol dsODN on ice. HEK293T cells were washed with PBS and trypsinized, then neutralized with media, spun down, and resuspended in PBS. Cells were counted and 4E5 cells per reaction were pelleted and resuspended in Lonza SF nucleofection reagent and supplement according to manufacturer’s instructions. 20 μL of cell solution was added to the RNP/dsODN mixture and then pipetted into a Lonza 16-well nucleocuvette.

Reactions were nucleofected using the CM-130 program. Warm media was added directly into the nucleocuvette after nucleofection, and cells were allowed to recover for 10 minutes at 37°C. After recovery, cell mixtures were transferred from the nucleocuvette to a 12-well plate containing pre-warmed media. Cells were harvested after 72 hours of growth, and GUIDE-seq library preparation was performed at GeneGoCell (San Diego, CA, USA).

For data processing, unique molecular identifiers (UMIs) were extracted from raw fastq files using umi-tools v1.1.4^54^. Adapters were trimmed off using cutadapt v.3.5^55^ and reads without correct adapters were discarded. Reads were aligned against the reference genome (hg38) using bwa mem v.0.7.17^56^ and deduplicated using umi-tools. Custom python scripts were used to identify sites that contained a dsODN, followed by re-alignment of these sites against the on-target, annotations of the genomic context of the detected sites, and generation of summary plots.

Sites were considered for inclusion if at least one read mapped to each side of the cleavage event, and the site was no more than 7 mismatches without DNA/RNA bulges or 6 mismatches and one DNA/RNA bulge (gap size = 1) away from the target sequence.

#### In vitro off-target nomination

Cas-OFFinder 3.0.0b3^39^ and the human reference genome (hg38) were used to identify reference sites meeting one or more of the following criteria for inclusion in a synthetic library of potential off-target sequences:

1. sites containing up to 6 mismatches in the PAM and guide sequence, with no DNA/RNA bulges

2. sites containing up to 4 mismatches, and 1 DNA/RNA bulge in the PAM and guide sequence, with a maximum bulge size of 1 base

3. sites containing up to 3 mismatches, and 1 DNA/RNA bulge in the PAM and guide sequence, with a maximum bulge size of 2 base

Sites associated with genetic variants were identified with Calitas^57^, using a list of genetic variants identified by the 1000 Genomes Project^58^ with a minimum global allele frequency of 1%. Two types of genetic variant sites were included in the library:

<u>1. Overlapping sites</u>: Sites detected among the reference sites, where the minimum number of mismatches plus gaps (MMG) for the variant-associated alignment is fewer than the minimum MMG for the reference alignment.

<u>2. De novo sites</u>: Sites not detected among the reference sites, where the variant-associated alignment meets any of the original three site inclusion criteria.

Negative control sites with no significant homology to the guide sequence were selected and added to the library. Target oligos were designed including the target sequence, 15 bp of flanking sequence on either side, and a unique 15 bp barcode for each site. Libraries were ordered from Agilent Technologies. The library was resuspended and amplified with 1x Kapa HotStart ReadyMix with primers including unique molecular identifiers (UMIs) and constant regions to generate cleavage ready libraries. A fraction of the purified cleavage ready libraries was prepared for Illumina sequencing to validate barcode representation within the library, and the remainder was retained for downstream assay execution.

XE apoenzyme and 2-fold molar excess guide RNA were combined and incubated at 37°C for 15 minutes. Cleavage reactions containing 30 ng of amplified library and 500nM pre-complexed RNP were prepared in 1X cleavage buffer (20 mM Tris pH 7.4, 150 mM NaCl, 5 mM MgCl2, 1 mM TCEP + 0.01% NP40). Reactions were incubated at 37 °C for two hours. Reactions were quenched by the addition of Proteinase K and DNA was isolated with paramagnetic beads.

Standard end repair and adapter ligation were carried out to prepare libraries for Illumina sequencing. DNA products of the expected size for cleaved and adapter-ligated substrates were isolated via size-selection with a PippinHT according to manufacturer instructions. The resulting products were PCR-amplified with Illumina-compatible adapters and sequenced on an Illumina NextSeq.

For data processing, paired-end FASTQ files were merged using flash v2.2.00^59^. Constant regions were trimmed off using cutadapt v.3.5^55^ and barcodes and unique molecular identifiers (UMIs) were recorded for each read. Custom python scripts were used to assign reads to library members by their barcode followed by alignment to the designed library sequence and identification of cleavage position. Reads with cleavage outside the expected window were discarded. Remaining reads were then deduplicated using the UMIs, and per-site enrichment values were calculated by dividing the representation of each barcode after cleavage by the representation in the uncleaved starting library. Sites were nominated as potential off-targets if their enrichment value was ≥1 or if their representation in the cleaved pool was ≥0.1% of the on-target site.

#### In silico prediction

Sites in the human genome with homology to the spacer of interest were identified using CasOffinder (Bae et al. 2014), with allowing up to 1 mismatch in the PAM sequence and the spacer sequence meeting any of the following criteria:

• ≤4 mismatches (MM) in the spacer sequence for sites in exonic regions (no bulges)

• ≤3 MM in the spacer sequence for sites in non-exonic regions (no bulges)

• ≤2 MM and 1 DNA/RNA bulge (length = 1) in the spacer sequence

#### rhAmpSeq

Primary human hepatocytes (PHH, Lonza lots HUM190131 and HUM181441) were plated at a density of 60K cells/well 24 hours prior to treatment with XE LNP at a dose ≥10x the EC90 as determined in prior potency experiments. Cells were maintained in sandwich culture for six days, at which point cells were digested with 1mg/ml collagenase/dispase for 20 minutes at 37C, harvested, and genomic DNA was extracted using Qiagen DNeasy Blood & Tissue Kit according to manufacturer’s instructions. Samples were prepared for sequencing using IDT’s rhAmpSeq CRISPR Library Kit and custom designed multiplexed rhAmpSeq panels according to manufacturer’s instructions. Off-target sites which were unsuitable for multiplexed analysis were assessed using a standard singleplex amplicon sequencing approach.

Multiplexed library pool sequencing data was analyzing using CRISPRessoPooled^60^. Each site in the designed pool was assigned a QC status based on the following criteria: a) the site has > 1000 reads assigned after the first sorting step of CRISPRessoPooled analysis, b) the site has > 95% alignment rate after the second alignment step of CRISPRessoPooled analysis, and c) the site has exactly 1 unique allele above 1% frequency in the untreated sample OR 2 unique alleles above 1% frequency within 5% frequency of each other with no unambiguous base calls (“N”). Sites that met all three criteria were selected for further analysis. We next used CRISPRessoCompare^60^ to compare the treated and untreated samples (Fisher’s exact test followed by Bonferroni correction with p-value cutoff 0.05). Any sites that showed significant deletions within the quantification window of the site (40 bp on either side of the cleavage site) were examined for final editing determination. Any single allele was deemed to be real editing if the treated sample contained at least 1 allele with a deletion 2 bp or larger spanning the cleavage site with a frequency above 0.5%, and if that same allele was not detected in the untreated sample. The on-target net indel rate was calculated as the average of two treated samples indel rate minus the untreated sample indel rate.

#### Statistical analysis and data visualization

Statistical tests were performed in GraphPad Prism unless otherwise noted. Schematic diagrams were generated using BioRender.

