## Supplementary Figures for "Comprehensive engineering of the CasX-Editor to generate a therapeutically optimized genome editor"

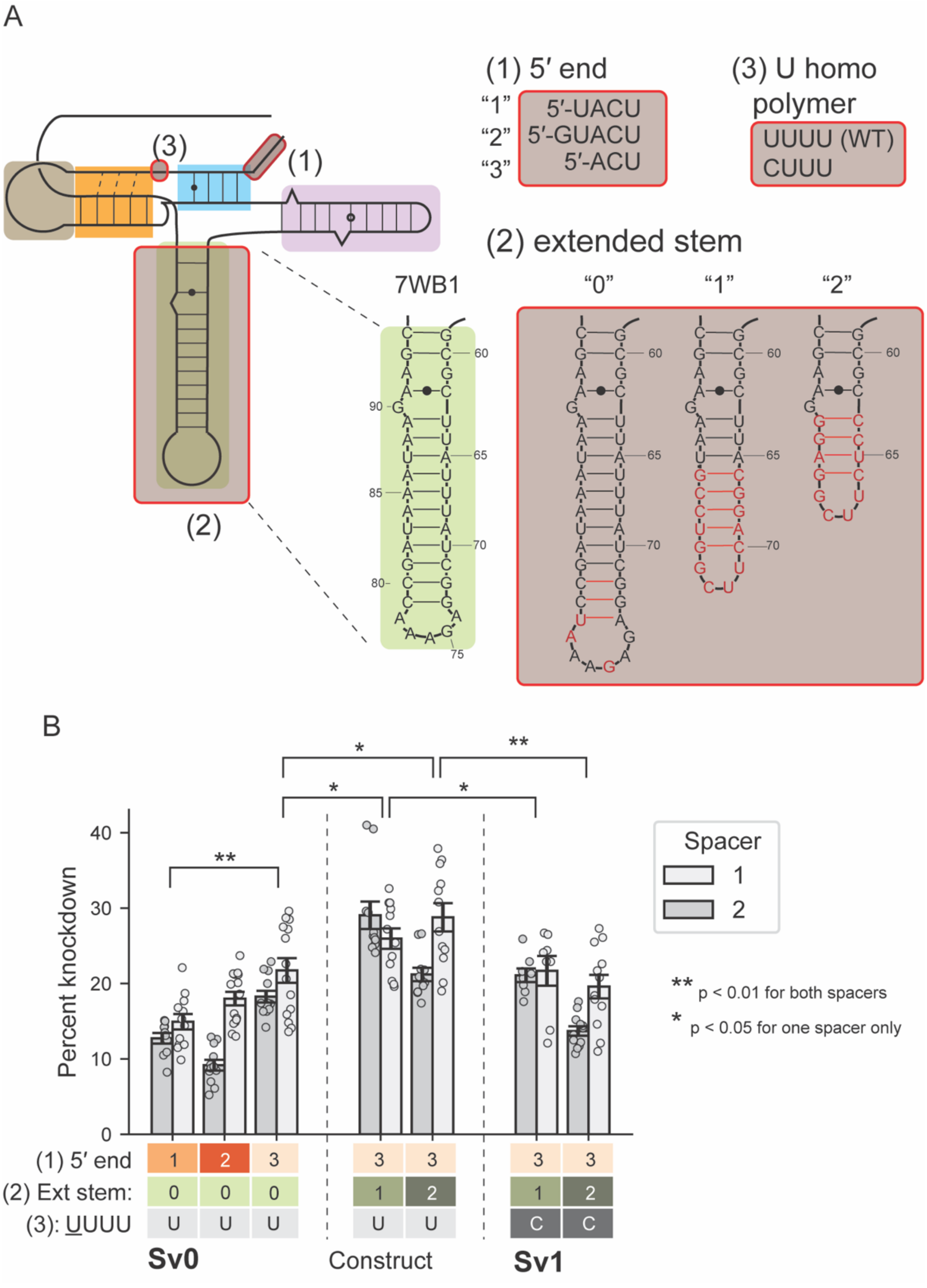


**Figure S1. Scaffold “Sv1” was identified from modifications targeting the 5′ end, extended stem, and a U-homopolymer region.**

(A) Sequence and predicted structure of the tested 5′ end modifications, the extended stem, and the homopolymer region. (B) Activity of the different scaffolds across two spacers targeting an endogenous cell surface receptor. Each point represents a replicate. Mean and standard deviation are shown.

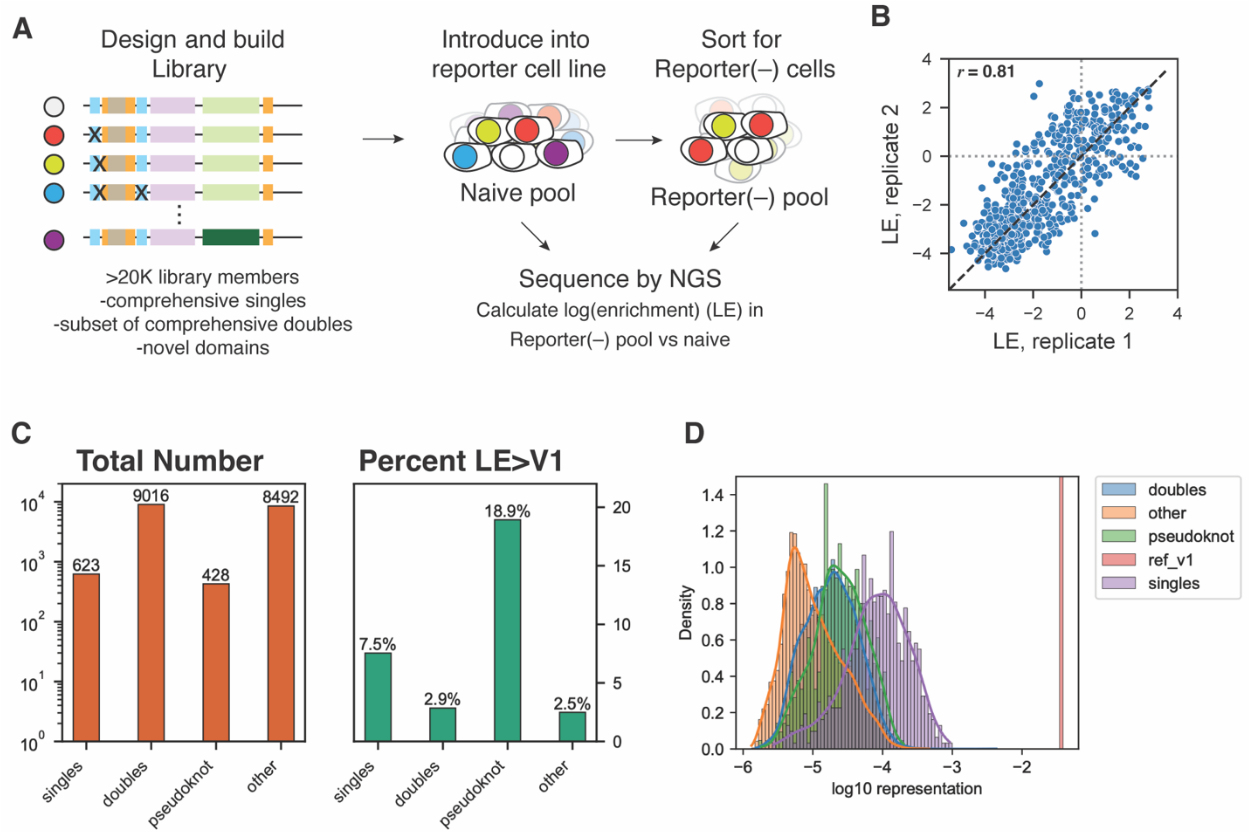


**Figure S2. Screening workflow identifies enriched single mutations.**
(A) Screening workflow from library design to calculation of log2 enrichment (LE) of variants in the reporter(-) pool versus naive pool. (B) LE ($log_2$ enrichment) for single mutations across two replicates demonstrates high reproducibility (r=0.81). (C) Total number of tested sequences (orange) and percent with positive LER (LE>V1; green) in the screen, separated by single mutations, double mutations, pseudoknot mutations, or other. (D) Representation of variants in the naive, unselected pool.

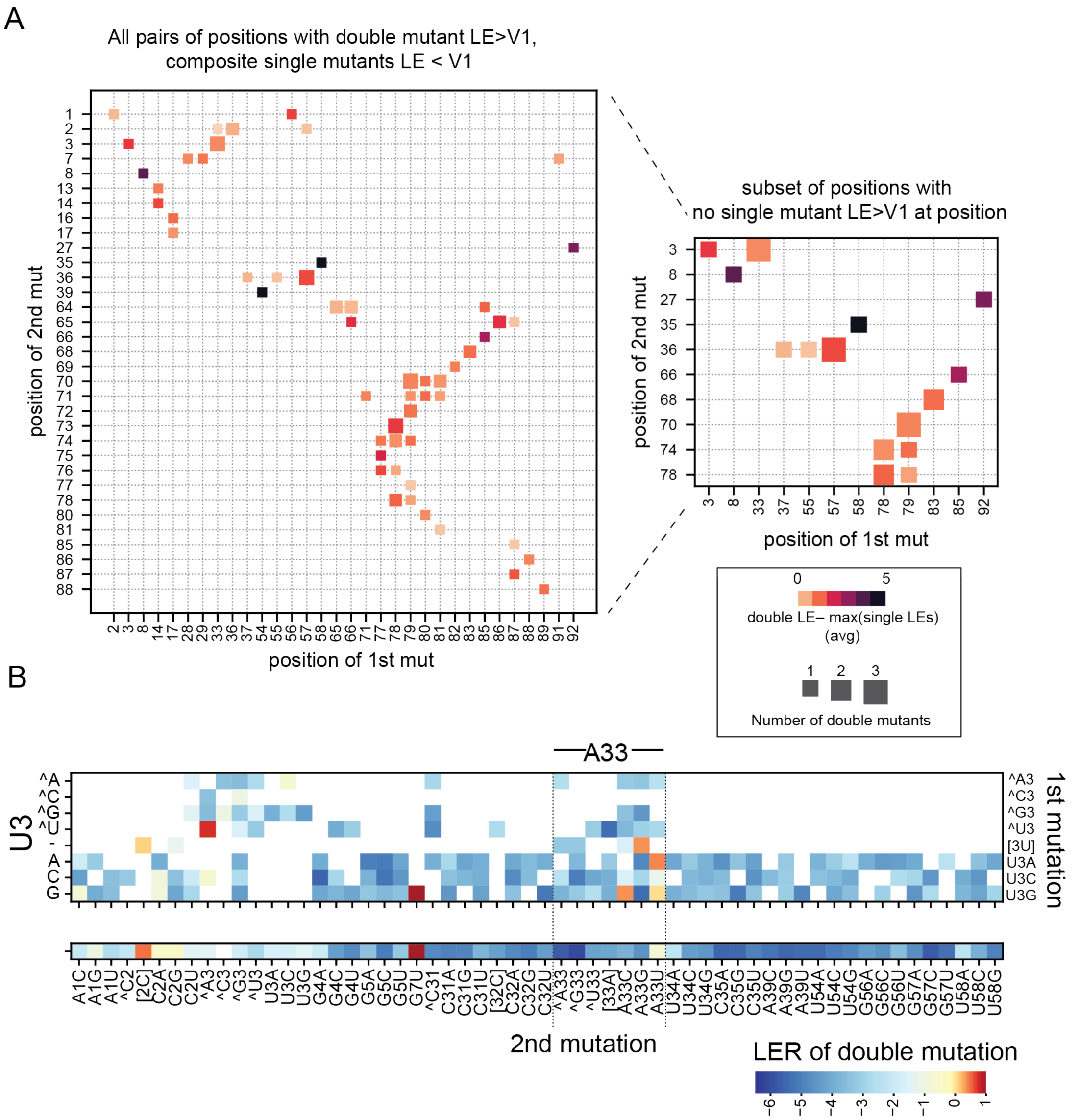


**Figure S3. Positive epistasis in the double mutations set.**

(A) Double mutants with LER >0 when neither composite single mutation had LER >0 were identified as demonstrating positive epistasis. The plots show the number of positive epistasis double mutations at that pair of positions (size of marker), and the average difference between the double mutation LE and the maximum of the composite singles LE. (Right) Same plot as left, but positive epistasis doubles are defined as those with LER > 0, with no single mutations at that pair of positions having LER > 0, which substantially reduced the number of mutations within this set. (B) All double mutations that address position 3. The U3-A33 double mutations show certain mutations resulting in canonical or non-canonical base pairs with positive LER (U3G,A33C; U3A,A33U; U3G,A33U; and [U3],A33G), while all single mutations at either site show negative enrichment. The enrichment for coordinated mutations that allow base pairing and deenrichment of mutations that would disrupt a predicted pair strongly suggests that U3 and A33 form a functionally significant pair not captured in published structures of CasX guides due to the use of a shortened 5′ end.

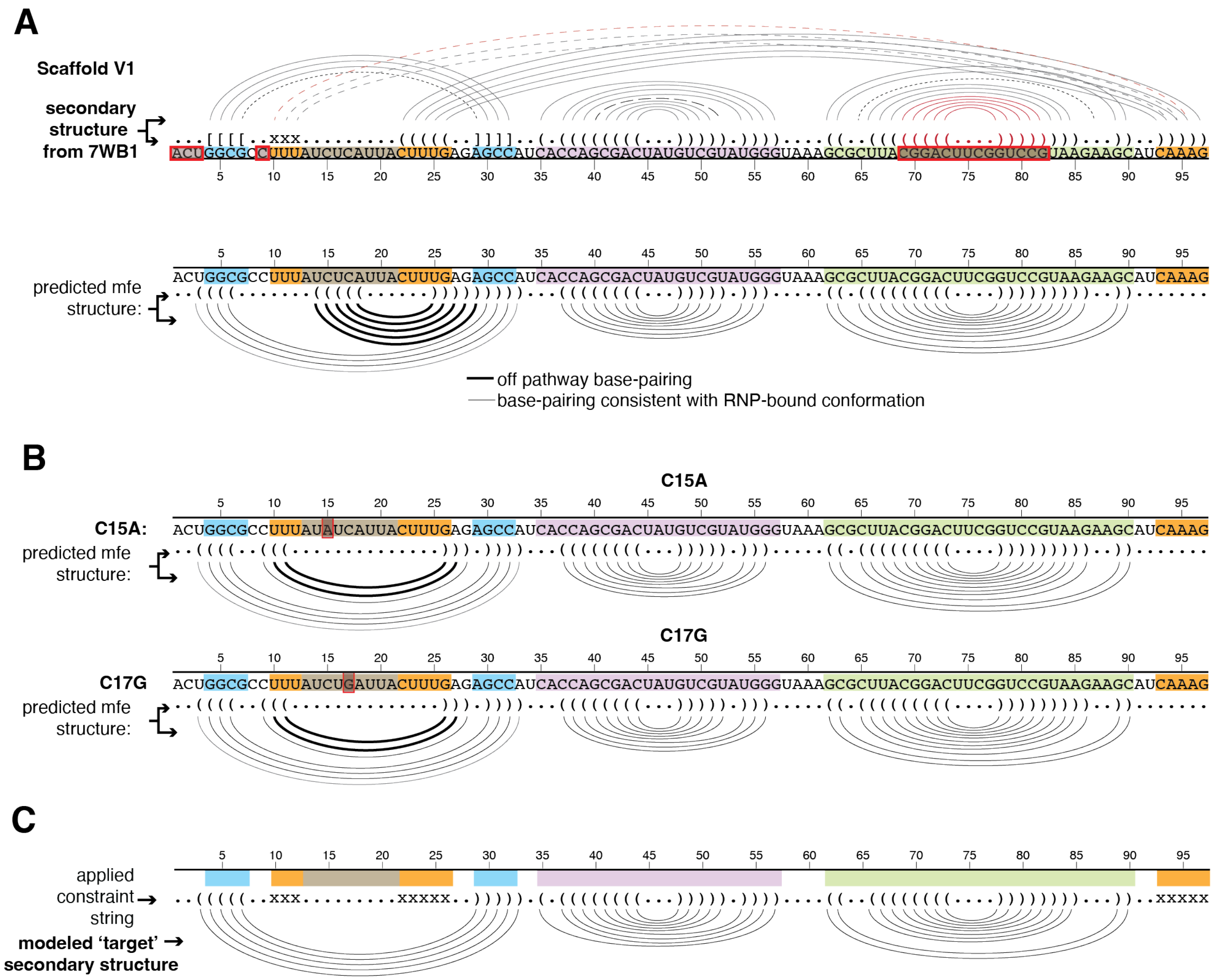


**Figure S4. Predicted structures of engineered scaffolds with and without constraints.** (A) Secondary structure of Sv1 as informed by the 7WB1 cryo-EM structure (top - lines and dot-bracket notation). Predicted mfe secondary structure for Sv1 identifies an off-target secondary structure with pairing in the triplex loop and pseudonkot stem. (B) Predicted minimum free energy (mfe) secondary structure for C15A and C17G partially resolves the off-target base pairing. (C) Modeled 'RNP-competent' secondary structure requires pseudoknot to be formed, and triplex bases to be unpaired. The secondary structure prediction algorithm employed here cannot predict pseudoknots or triplex energetics, thus this simplification was used to predict the states competent for triplex folding.

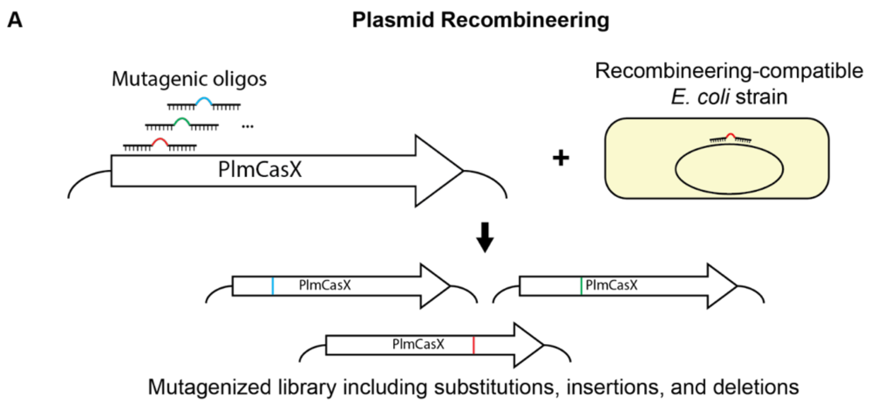


**Figure S5. CasX variant library generation.**
(A) Schematic of plasmid recombineering workflow. A library of mutagenic oligos encoding desired mutations (all possible single insertions, deletions, and substitutions) of PlmCasX are electroporated into a recombineering-compatible *E. coli* strain. The mutagenic oligos are incorporated as the plasmid is replicated, generating a plasmid library.

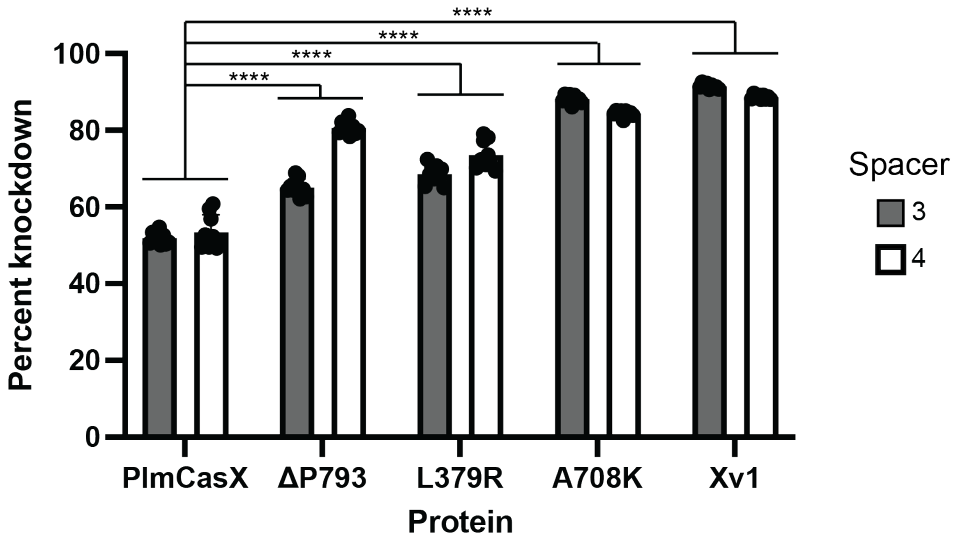


**Figure S6. Plasmid transfection of single-mutation protein variants.**
Plasmids encoding single-mutation variants of PlmCasX and scaffold Xv6 guides with B2M-targeting spacers 3 and 4 were transfected into HEK293 cells. The three tested single mutations improved activity over PlmCasX with both spacers. p-values relative to PlmCasX for the matching spacer are shown (one-way ANOVA, Dunnett's multiple comparison test). **** = p<0.0001.

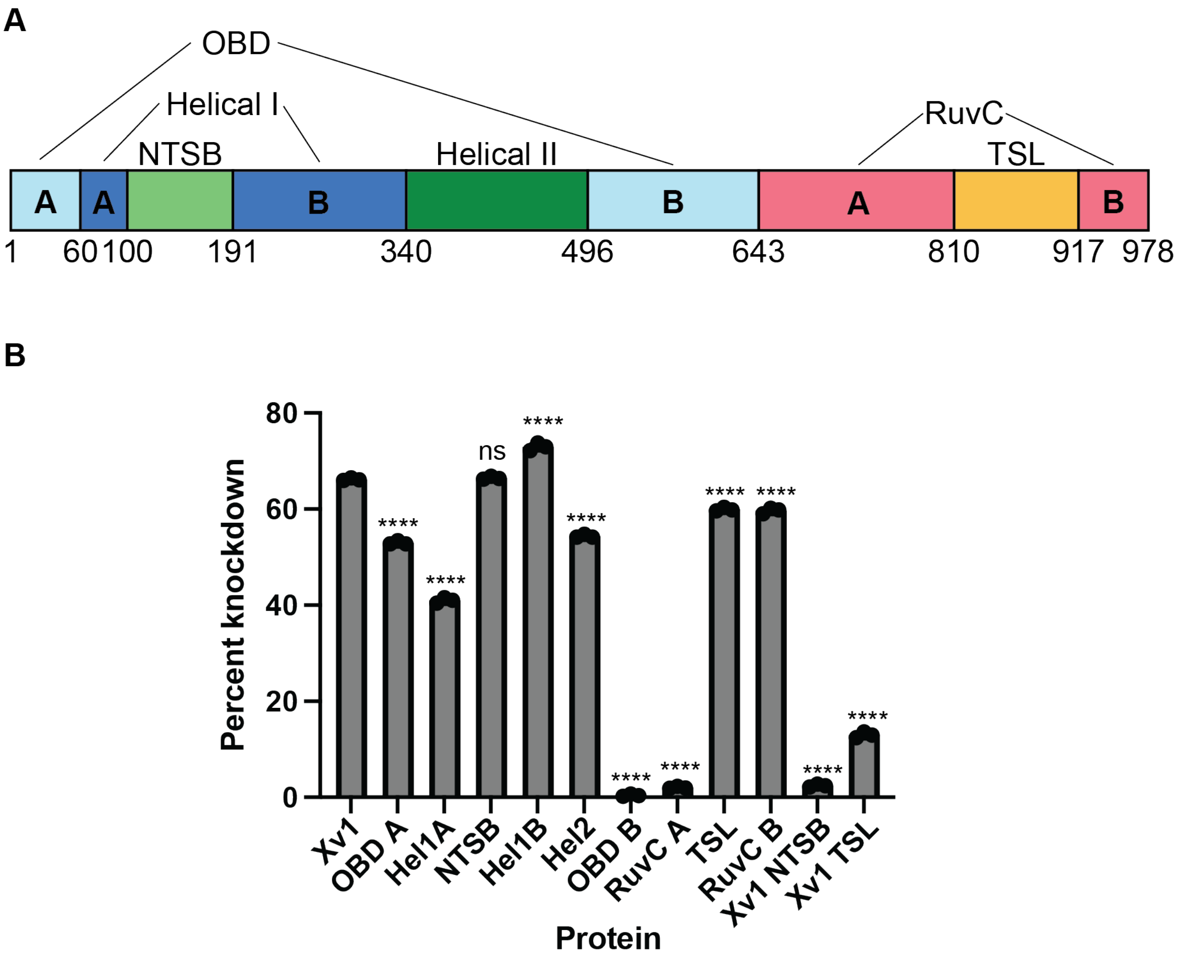


**Figure S7. Domain exchanges have variable effects on potency.**
(A) Domain annotations and boundaries on Xv1 used to guide domain exchanges. Domains based on Tsuchida et al., except that the bridge helix was considered part of RuvC A. (B) Percentage of reporter-negative cells when treated with plasmid coding for an XE variant and a reporter-targeting guide. For each protein, the indicated domain from DpbCasX replaced the corresponding domain in Xv1, except for Xv1 NTSB and Xv1 TSL, where the indicated domain from Xv1 replaced the corresponding domain in a DpbCasX base molecule. Mean, standard deviation, and individual measurements for three replicates are shown. p-values relative to Xv1 are shown for each bar (one-way ANOVA, Dunnett's multiple comparison test). **** = $p<0.0001$, ns = not significant ($p>0.05$).

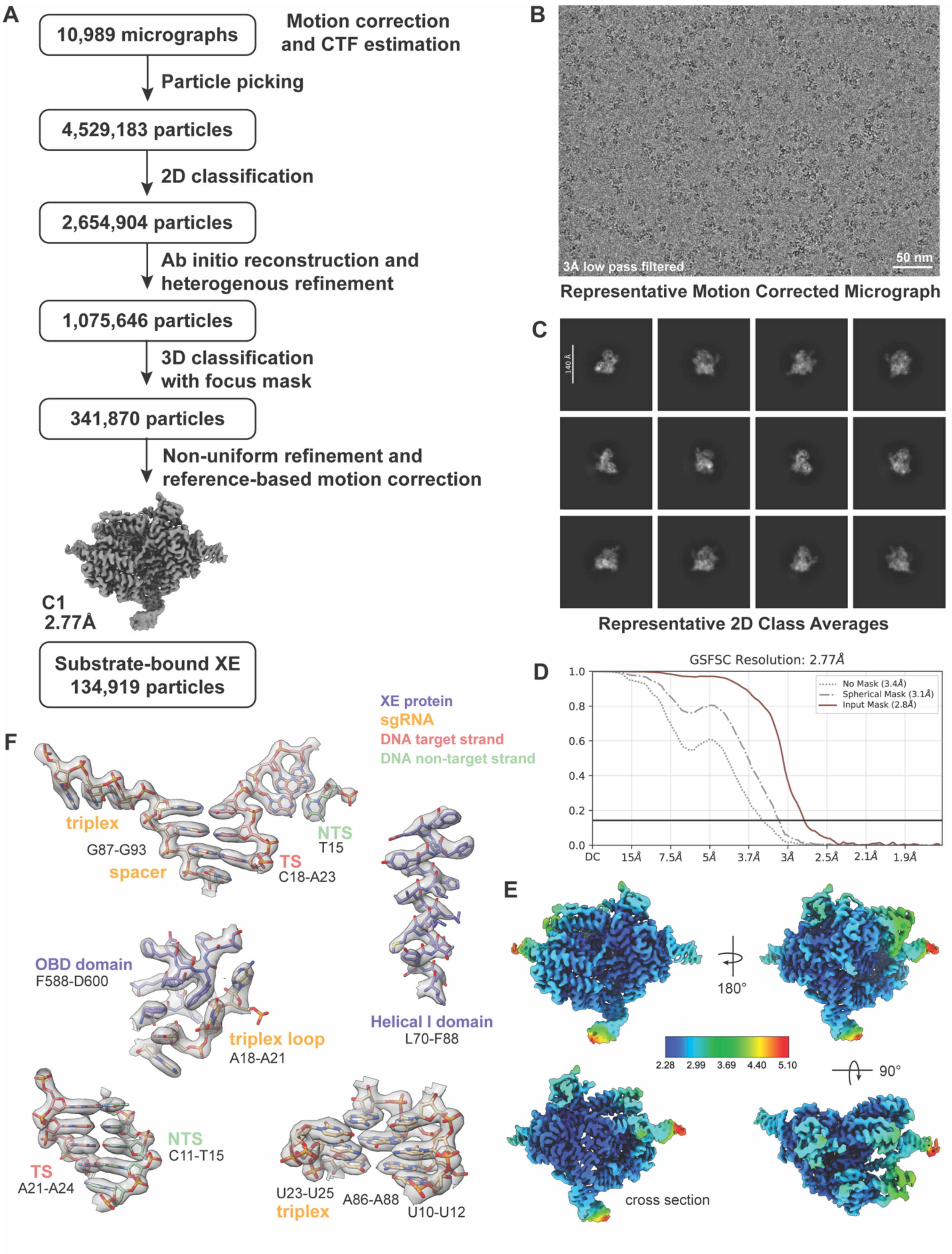


**Figure S8. Cryo-EM data analysis for substrate-engaged XE reconstruction.**

(A) Overall data processing workflow for the reconstruction of XE engaging its target DNA substrate. (B-C) Representative micrograph with scale bar (50 nm) and 2D classes. (D) Gold-standard Fourier Shall Correlation (GS-FSC) curve. (E) Local resolution. Scale bar in Å. (F) Representative examples for the cryo-EM map and the corresponding modeled regions. Map has transparent surface; model is shown with sticks and heteroatom coloring (red for oxygen, orange for phosphorus, yellow for sulfur, blue for nitrogen).

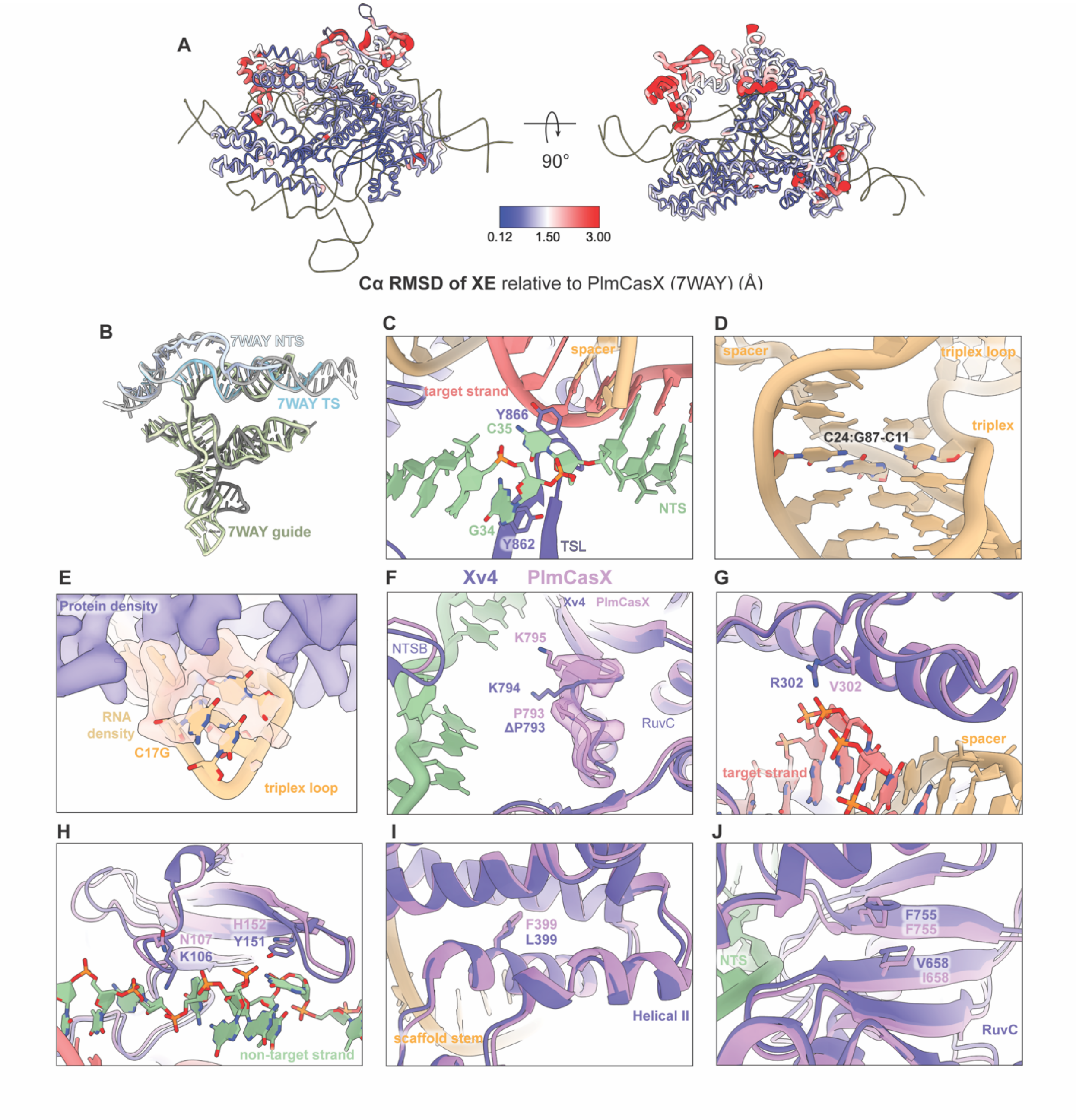


**Figure S9. Structural impact of mutations in XE**.

(A) XE enzyme model colored by the Cα RMSD relative to PlmCasX (PDB: 7WAY) (B) Overlay of XE guide RNA and DNA substrate with that of PlmCasX (PDB: 7WAY) (C) Detailed view of aromatic residues interacting with non-target strand at PAM-distal end of R-loop. (D) Detailed view of coordinated triplex mutation. The expected C-G:C triple is formed and shown in heteroatom coloring. (E) Cryo-EM density (shown as transparent surface) and model for triplex loop, including C17G mutation. C17G shows poor density and no interactions with protein. U14-G17 model shown in heteroatom coloring. (F) Local impact of ΔP793 is shown. PlmCasX model (PDB: 7WAY) is shown in light purple, with corresponding cryo-EM density shown as transparent surface for the residues around P793. XE model is shown in dark purple. Same coloring is used for F-J. (G) Helical I domain exchange introduces a V302R mutation that likely improves target strand binding (target strand shown in pink). (H) NTSB domain exchange introduces N107N and H152Y mutations that likely improve non-target strand binding (non-target strand shown in green). (I) F399L does not notably alter local structure. (J) V658I mutation appears to induce F755 to adopt a different rotamer but does not otherwise notably affect local packing.

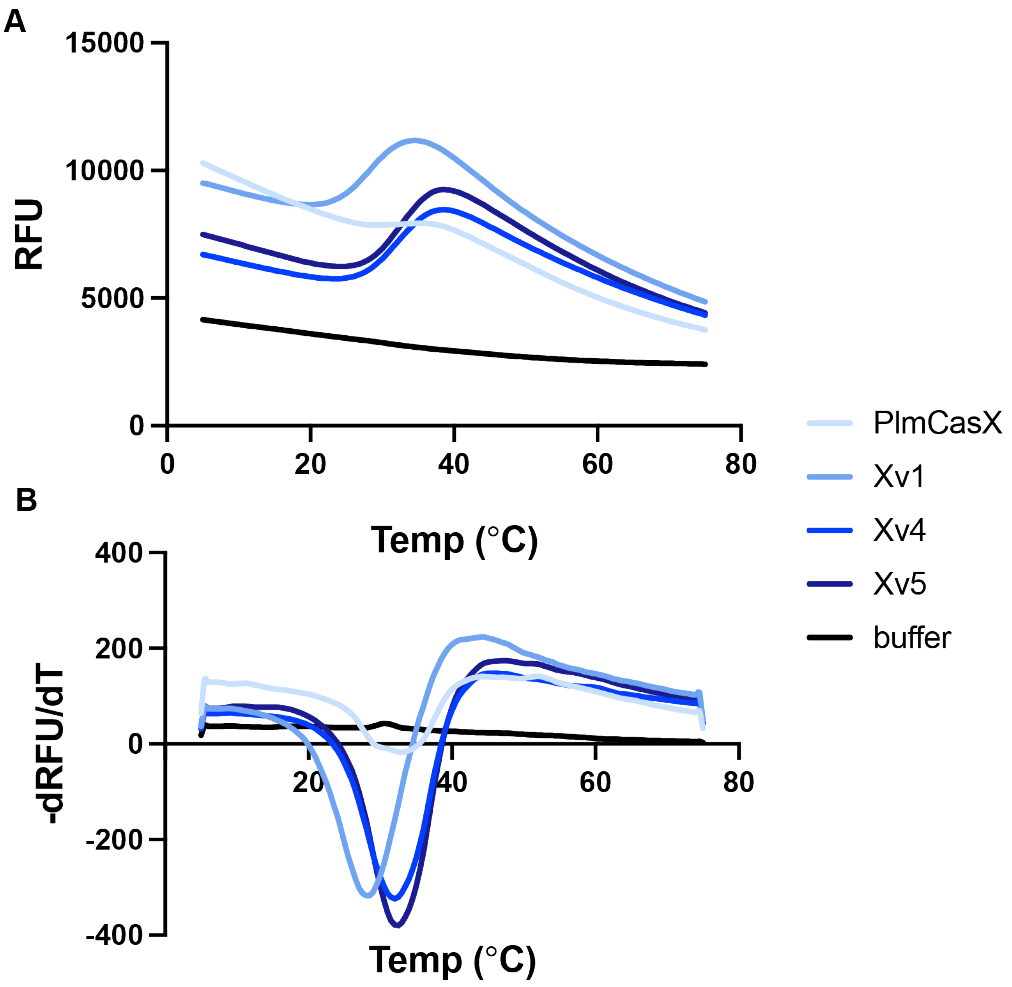


**Figure S10. Thermal shift traces.**

Fluorescent thermal shift assay of apoenzyme CasX variants shown as RFU values (A) and the change in RFU in response to change in temperature (B). PlmCasX did not show a clear transition in the range of the other variants, preventing confident calculation of Tm. PlmCasX was observed to precipitate when kept on ice as well as at room temperature or above, suggesting that it might be poorly folded even at low temperature in the tested buffers. Exposed hydrophobic regions would explain both the tendency to aggregate and lack of clear fluorescence increase in the attempted melt curve.

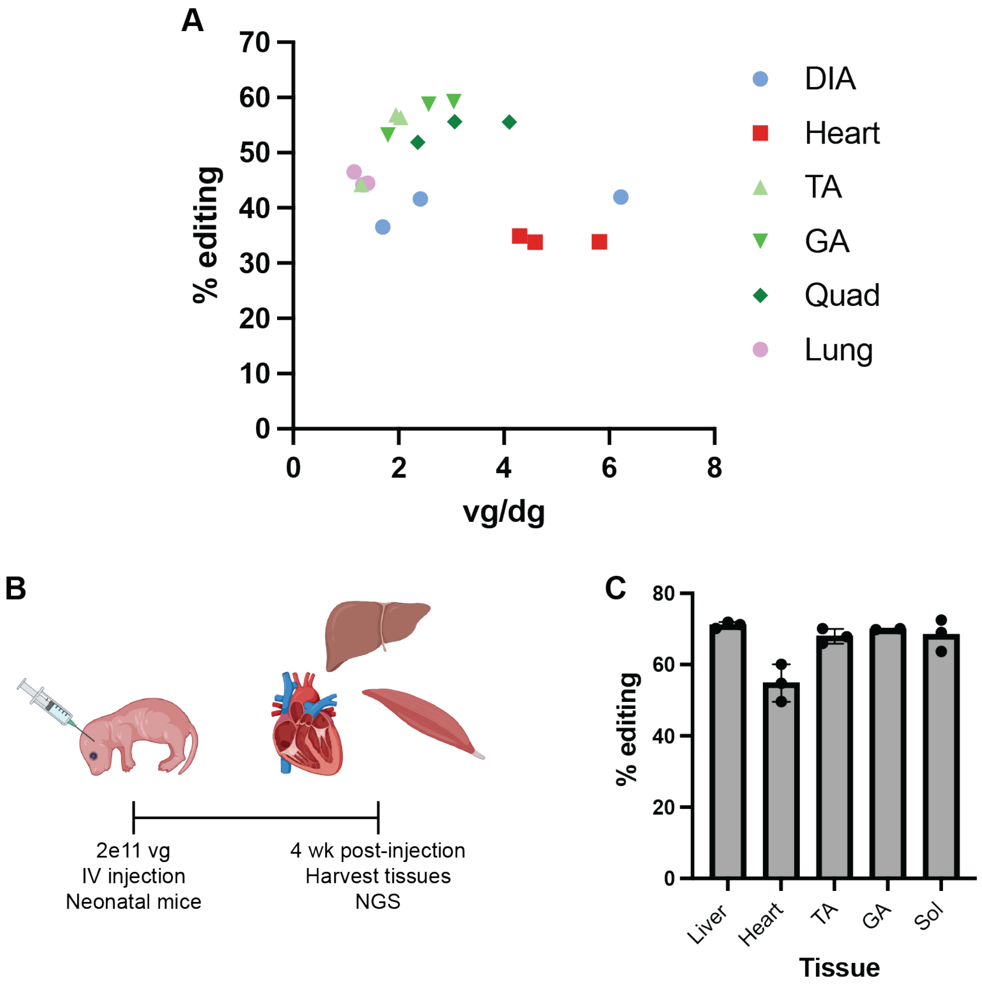


**Figure S11. AAV-mediated editing of muscle in mice.**

(A) Editing rate (measured by NGS) plotted vs viral genomes per diploid genomes (measured by ddPCR) for different tissues for adult mouse experiment presented in Fig 4C-E. Each point represents data from a tissue sample taken from a single mouse. (B) Experimental design of systemic AAV delivery in neonatal mice. (C) Editing at *ROSA* locus measured by NGS. TA = tibialis anterior. GA = gastrocnemius. Sol = soleus. N=3 for liver, heart, TA, and Sol. N=2 for GA.

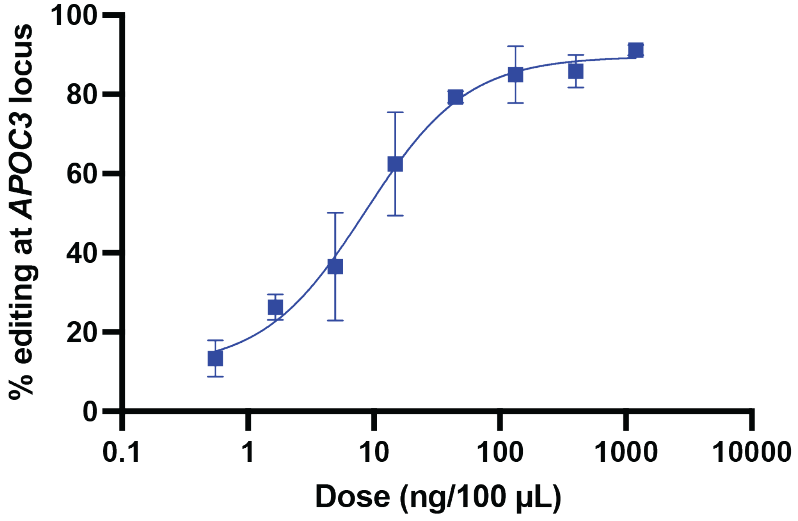


**Figure S12. *APOC3* editing in primary human hepatocytes.**
Editing at the *APOC3* locus as measured by NGS in primary human hepatocytes following administration of LNP with Xv5 mRNA and Sv7 guide targeting *APOC3* at the indicated dose. Mean and standard deviation of three experimental replicates is shown, along with a four-parameter logistic model fit.

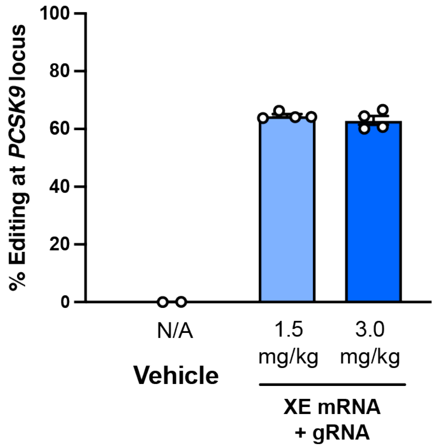


**Figure S13. Saturation of on-target editing of the *PCSK9* locus in mice.**
A humanized transgenic mouse model (KIKO) was treated with a single intravenous dose of LNP-formulated Xv5 mRNA and *PCSK9*-targeting Sv7 gRNA. Seven days post-treatment, editing of the *hPCSK9* locus was assessed. N=2 for saline control; N=4 for 1.5 mg/kg and 3.0 mg/kg dose groups. Points represent individual animals; bars show on mean ± s.e.m.

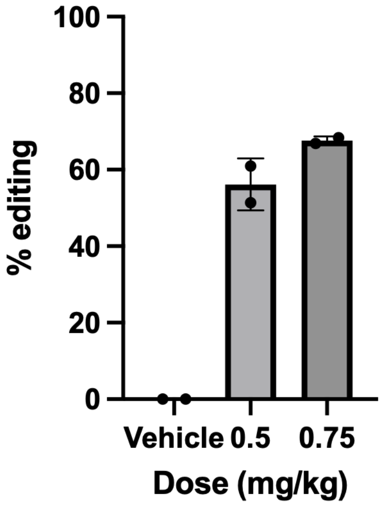


**Figure S14. Editing of the *PCSK9* locus in non-human primates at sub-saturating doses.** Editing in liver tissue of cynomolgus macaques following a single intravenous infusion of *PCSK9*-targeting XE LNP. Vehicle control animals received PBS. Liver punch biopsies were collected at Day 15. N=2 for each dose group. Individual points represent animals; bars show mean ± standard deviation.

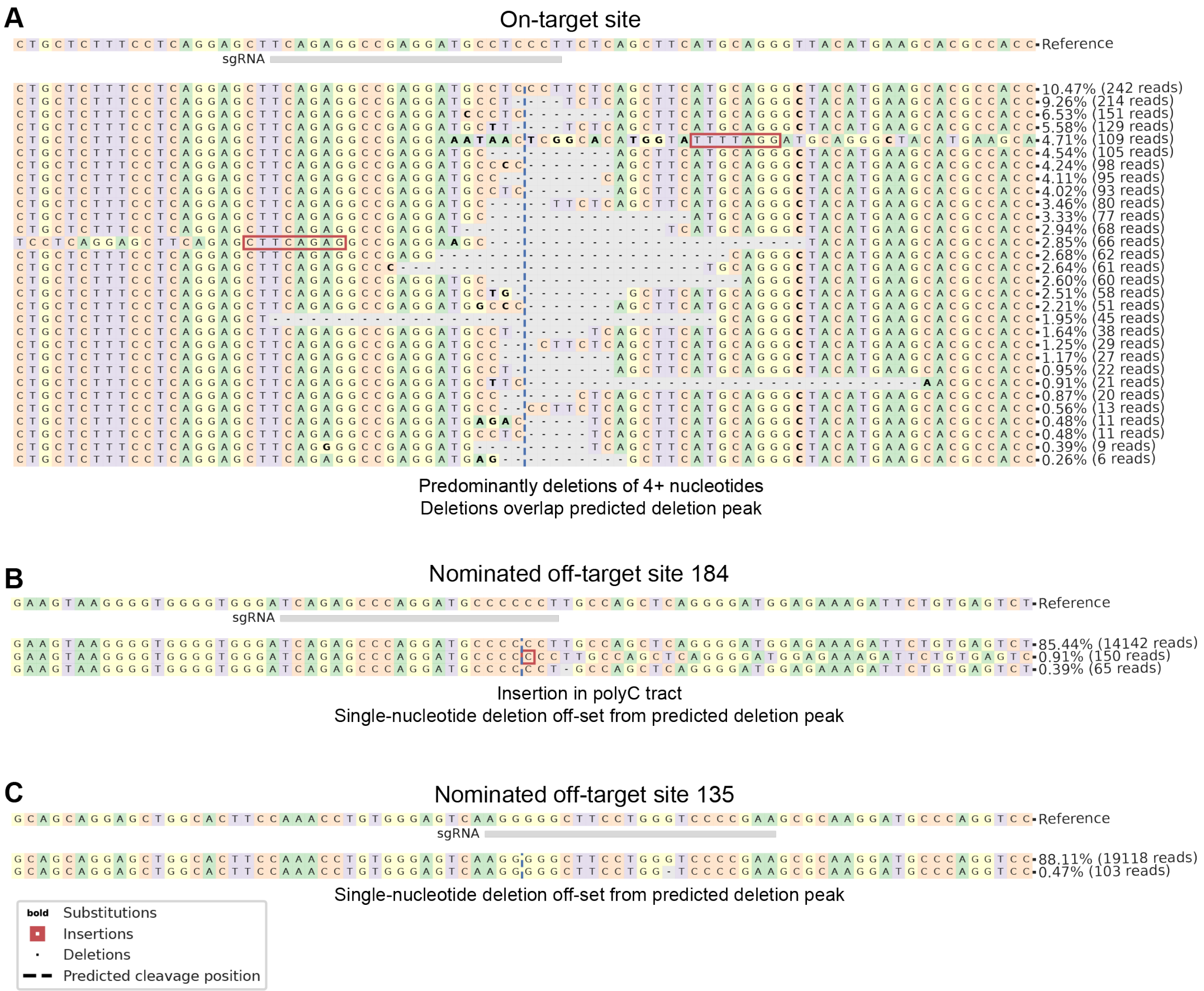


**Figure S15. Analysis of nominated off-target sites for APOC3.**

(A) Reads at the *APOC3* target site with indels. The stereotypical peak deletion site is plotted. Deletions are nearly all 4 nt or greater, and all overlap or flank the predicted deletion peak. (B) Indel-containing reads at a potential off-target site. The most common indel allele shows a single-nucleotide insertion, which is not a common outcome of editing by Type V enzymes. The insertion is also within a polyC tract, increasing the likelihood of sequencing error. The next most common allele is a single deletion offset from the predicted deletion peak, which is also atypical of XE edits. (C) Indel containing reads at a potential off-target site. A single allele is identified with a single-nucleotide deletion, >10 nt offset from the predicted deletion peak. This profile is inconsistent with XE edits.

**Table S1 – Non-human primate editing rates of clinical CRISPR-based gene editors**

| **Editor:** | XE | XE | ABE8.8[1] | SpCas9[2] | SpCas9[3] | SpCas9[4] |
|---|---|---|---|---|---|---|
| **Target:** | *PCSK9* | *APOC3* | *PCSK9* | *Lp(a)* | *ANGPTL3*** | *TTR**** |
| **Dose (mg/kg)** | **Editing rate** | | | | | |
| 0.5 | **49.9%, 56.2%*** | **64.6%** | | 10% | 45% | |
| 0.75 | **67.6%** | | 46% | | | |
| 1.0 | | **77.1%** | | | | |
| 1.5 | **76.1%** | | 70% | 45% | 66% | 48% |
| 2.0 | | **81.1%** | | | 68% | |
| 3.0 | **77.6%** | | | 65% | | 76% |

*Averages from two separate experiments

**Values estimated from graph

***Phenotypic knockdown scaled to expected fraction of edited cells, assuming maximum targetable fraction of liver cells is 80% and a 1:1 relationship between editing rate and phenotypic knockdown
